# Bacteria primed by antimicrobial peptides develop tolerance and persist

**DOI:** 10.1101/802207

**Authors:** Alexandro Rodríguez-Rojas, Desiree Y. Baeder, Paul Johnston, Roland R. Regoes, Jens Rolff

## Abstract

Antimicrobial peptides (AMPs) are key components of innate immune defenses. Because of the antibiotic crisis, AMPs have also come into focus as new drugs. Here, we explore whether prior exposure to sublethal doses of AMPs increases bacterial survival and abets the evolution of resistance. We show that *Escherichia coli* primed by sublethal doses of AMPs develop tolerance and increase persistence by producing curli or colanic acid. We develop a population dynamic model that predicts that priming delays the clearance of infections and fuels the evolution of resistance. The effects we describe should apply to many AMPs and other drugs that target the cell surface. The optimal strategy to tackle tolerant or persistent cells requires high concentrations of AMPs and fast and long-lasting expression. Our findings also offer a new understanding of non-inherited drug resistance as an adaptive response and could lead to measures that slow the evolution of resistance.

## INTRODUCTION

Antimicrobial peptides - short, usually cationic peptides - are key effectors of innate immune defences of all multicellular life **(*1*)** and are also important players at the host microbiota interface **(*2*, *3*)**. Because of their evolutionary success and diversity, AMPs are considered as new antimicrobial drugs to alleviate the antibiotic resistance crisis **(*4*)** with currently more than two dozen under clinical trial **(*5*)**. *Bona fide* genetic resistance in AMPs has been studied **(*6*, *7*)**, but not to the extent of antibiotic resistance. Resistance against AMPs evolves usually with a low probability and the levels of resistance are not as high as against antibiotics **(*8*, *9*)**.

Notably, non-inherited resistance **(*10*)**, the ability of bacteria to survive lethal concentrations of antimicrobials without a genetically encoded resistance mechanism, has hardly been studied for AMPs. One of the very rare studies that has addressed this in a natural host-microbe interaction is the example of the bobtail squid and its symbiont *Vibrio fischeri*. Here, phenotypic resistance is elicited by a low pH and primes the *Vibrios* to colonize the light-emitting organ of the squid **(*11*)** in the presence of high concentrations of AMPs. Microbes have evolved adaptive physiological alterations to predictable environmental changes **(*12*, *13*)**. This has been studied in the context of available carbon sources during gut passage **(*12*)**, and a meta-analysis found that priming, the phenotypic response to a low level stressor, provides a fitness benefit for microbes against a variety of stressors including pH, temperature and oxidative stress **(*14*)**.

Here we study if the previous encounter with sub-lethal dosages of AMPs induces tolerance or persistence. Such low concentrations are common. For example, after infection the induction of AMPs results in sublethal concentrations before killing concentrations are reached. This usually takes a few hours **(*15*)**. Such pharmacokinetic profiles are mirrored in many antimicrobial drug treatments. During medical application of antibiotics, the pharmacokinetics start at zero and the killing concentrations are building up over time.

Resistance can be induced by sublethal levels of antimicrobials **(*16*)**. Non-inherited resistance, resulting in either drug tolerance or persister cell formation (see Fig. S 1, **(*17*)** has been shown to be of great importance to understand antibiotic resistance **(*10*, *16***–** *18*)**. It can also facilitate *bona fide* resistance evolution against antibiotics **(*18*)**. We adopt these findings and concepts from antibiotic research to understand resistance against AMPs, an insight which should equally inform our understanding of host-microbe interactions as well as resistance evolution against AMPs as drugs. It is noteworthy that AMPs differ significantly from conventional antibiotics in several aspects including their pharmacodynamics, resulting in narrower mutant selection windows **(*9*)**. Also, the speed at which they kill cells is fast, as they kill within minutes **(*19*)** rather than within hours, as it is the case for antibiotics **(*20*)**.

In this study, we specifically investigate whether prior exposure to sublethal doses of AMPs increases bacterial survival via either tolerance or persistence, and the risk of resistance evolution. We use two antimicrobial peptides that are well characterised as a case study. Melittin is a 26 amino acid residue linear peptide from the honeybee, which kills bacterial cells by permeabilization of the membrane **(*21*)**. Melittin is also active against eukaryotic parasites such as Leishmania and cancer cells. The other AMP used here, pexiganan, is the first eukaryotic AMP developed as a drug, a synthetic 22 amino acid residue peptide closely related to magainin from the African clawed frog **(*22*)**. It shows a broad activity against both, gram^+^ and gram^−^ bacteria and kills by forming toroidal pores **(*23*)**. We combine *in vitro* experiments with a modelling approach to study priming and resistance emergence. We find that a sublethal dose of certain antimicrobial peptides can induce increased tolerance and/or persistence in bacteria and hence prime **(*24*)** them for the exposure to a subsequent lethal dose. We identify candidate underlying molecular mechanisms and capture the population dynamics by adapting a classic mathematical model of persistence **(*25*)**. With computer simulations, we then predict that increasing tolerance and persistence will have a positive effect on bacterial survival and the emergence of resistance.

## RESULTS

### Primed cells are more resistant

We primed *E. coli* K-12 in vitro by exposing them to a fraction of the minimal inhibitory concentration of an AMP (0.1xMIC, table S1). Subsequently, we exposed the primed bacterial populations to lethal concentrations of the respective AMPs (10 × MIC, table S1) and then monitored bacterial survival over time. We found that the priming treatment resulted in much higher *E. coli* survival (Fig. 1 A, B).

**Figure 1:**
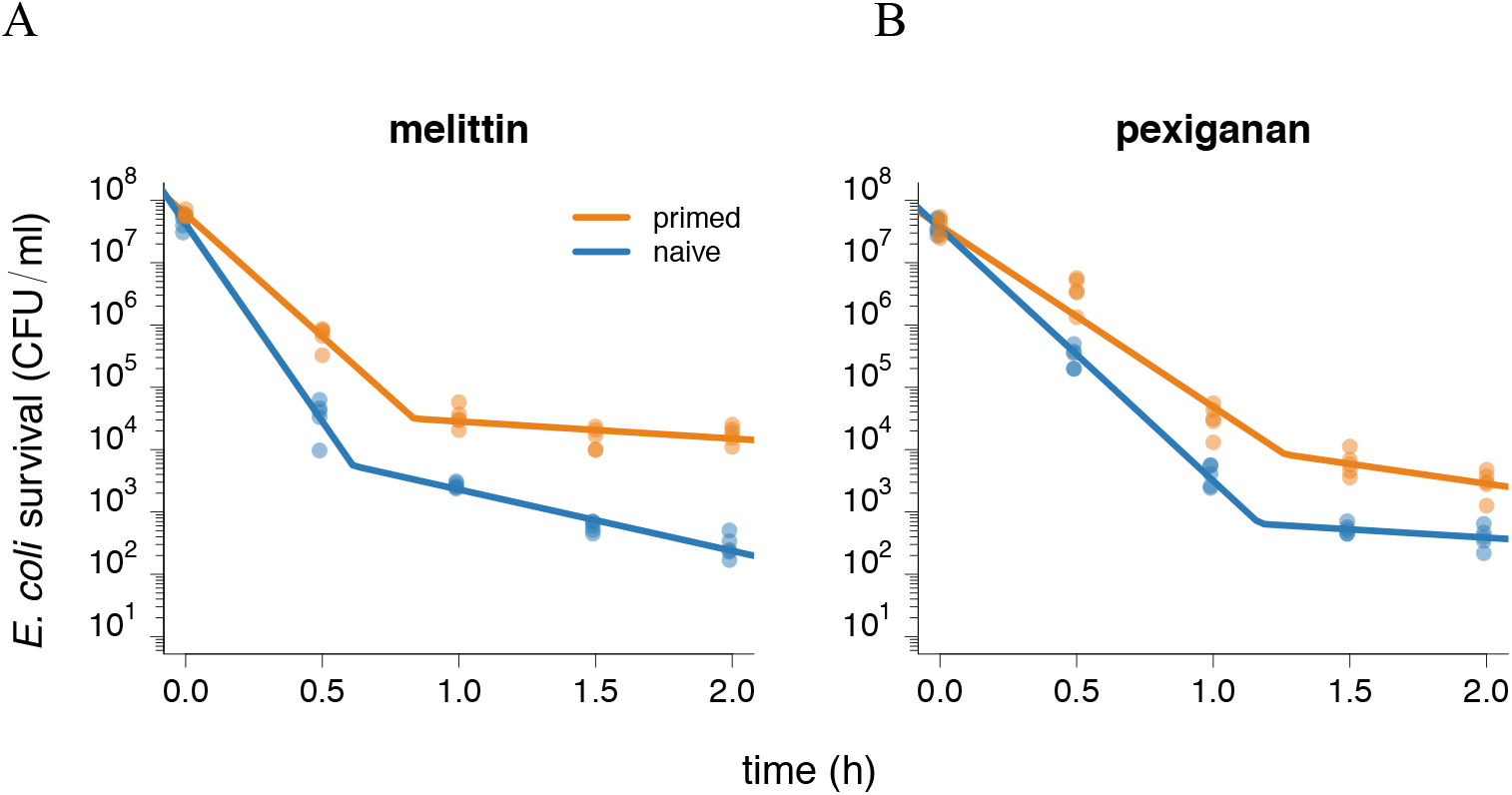
Bacterial tolerance and persistence determine the shape of time-kill curves. Time-kill experiments with *E. coli* K-12, primed (orange) and naïve (blue) bacteria were exposed to 10xMIC of (A) melittin and (B) pexiganan. At each timepoint indicated, we measured the bacterial population size 5 times. We tested with an ANOVA by means of contrasts if priming influenced tolerance and persistence. For both antimicrobials, priming significantly increased the slope of the first phase, the measure of tolerance, and the bacterial level in the second phase, the measure of persistence, (significance level: p < 0.05). We corrected for multiple testing with the Bonferroni-method. The line in the plots indicates the best fit of a biphasic function (table S2), on which our statistical analysis is based.

### Killing is biphasic

The decline of the time kill curves is biphasic suggesting two subpopulations (*25*–*28*). We excluded that deviations from monophasic decline arise because of decreasing antimicrobial concentrations over time (Fig. S 2) and fitted the time-kill curves to a biphasic linear function. For both AMPs, bacterial populations declined faster during the first than the second phase (Fig. 1, Table S 2). Tolerance, the decline of bacterial populations in the first phase (*17*), was significantly higher in primed than in naïve bacteria for both AMPs. Primed bacteria showed higher survival in the second phase, indicating higher numbers of persisters. The change in population size in the second phase, however, was not significantly different between primed and naïve populations, indicating that the population dynamics as such in the second phase are not influenced by priming (Fig. 1). In short, priming with AMPs allow bacteria to survive better by increasing both bacterial tolerance and persistence.

### Priming is mediated by either curli or colanic acid

To understand how priming leads to tolerance and persistence, we used RNAseq of cells exposed to priming concentrations of AMPs (Fig. 2 A, B). Exposure to sublethal concentrations of pexiganan (0.1xMIC, as above) induced colanic acid synthesis (Fig. 3 A). Colanic acid capsules have been shown to protect against AMPs and antibiotics **(*29*)**. The removal of an essential gene for colanic acid production completely abolished the priming effect by pexiganan (Fig 3 B). By phase contrast imaging, we observed the formation of a characteristic colanic acid capsule in pexiganan-primed but not in naïve cells (Fig. 3 C). The priming response was homogenous, with all observed cells producing a colanic acid capsule under priming conditions. Exposure to a sublethal concentration of melittin (0.1xMIC, as above) induced up-regulation of curli fimbriae (Fig 3 D). Curli is an important virulence factor **(*30*)** and a component of extra-cellular matrix that protects against AMPs **(*31*)**. A curli-deficient mutant showed a decrease of the priming effect induced by melittin (Fig. 3 B). For melittin-primed bacteria, curli induction was documented with a specific curli binding chemical (Fig. 3D). Both AMPs also induced significant overlap in gene expression related to osmotic shock (Fig. 2).

**Figure 2.**
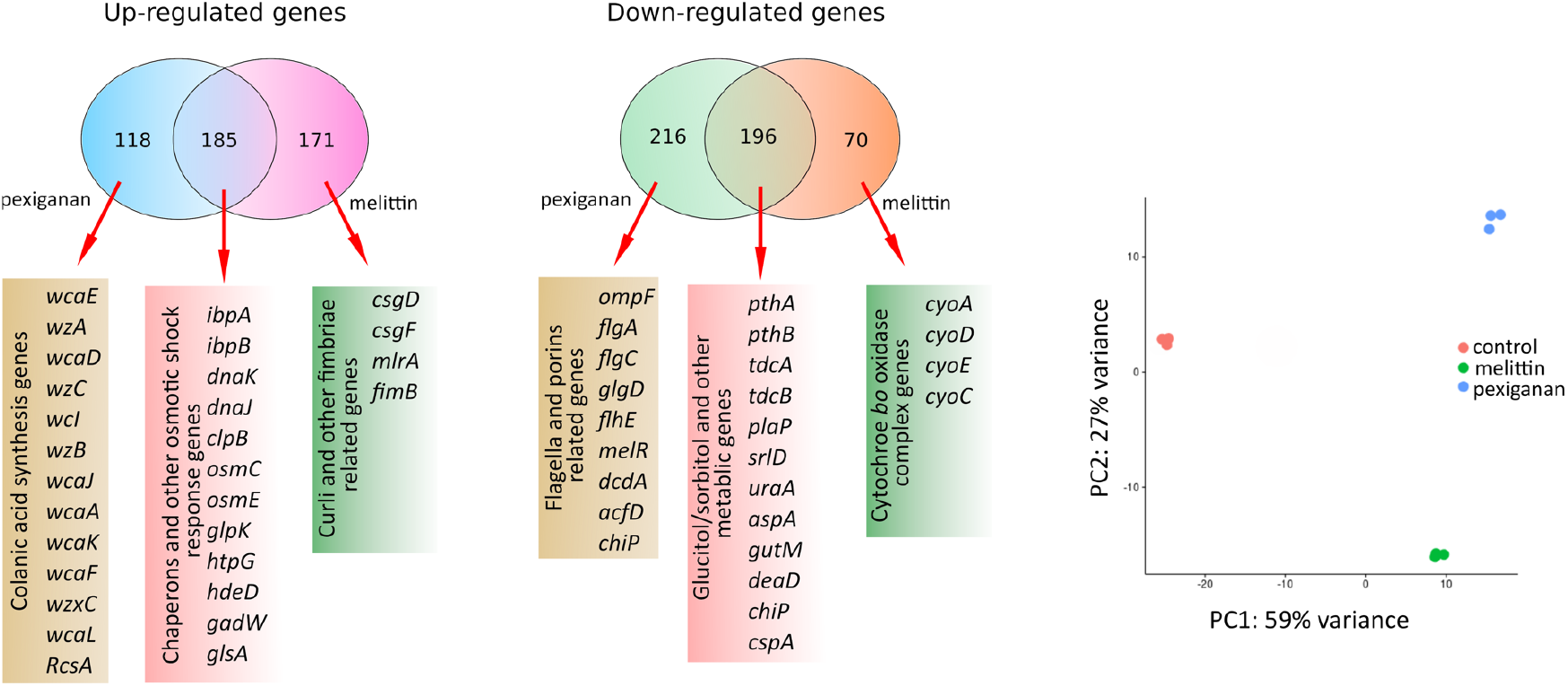
Gene expression in primed *E. coli*. (A) Venn diagrams showing specific and overlapping responses of *E. coli* MG1655 to priming concentrations of melittin and pexiganan (B) Principal component 1 separates the control from the peptide priming, PC 2 separates the melittin induced response from the pexiganan response.

**Figure 3.**
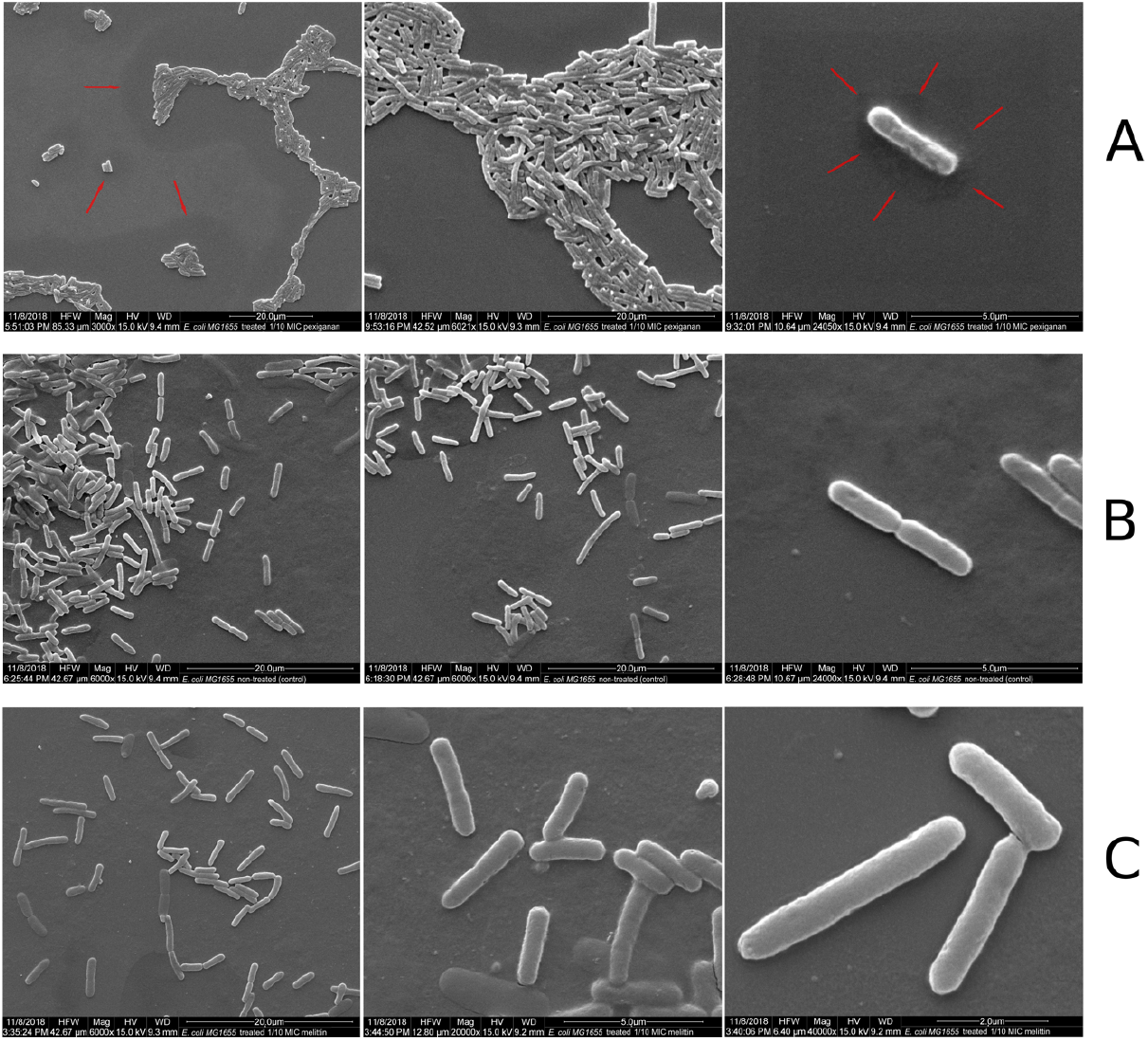
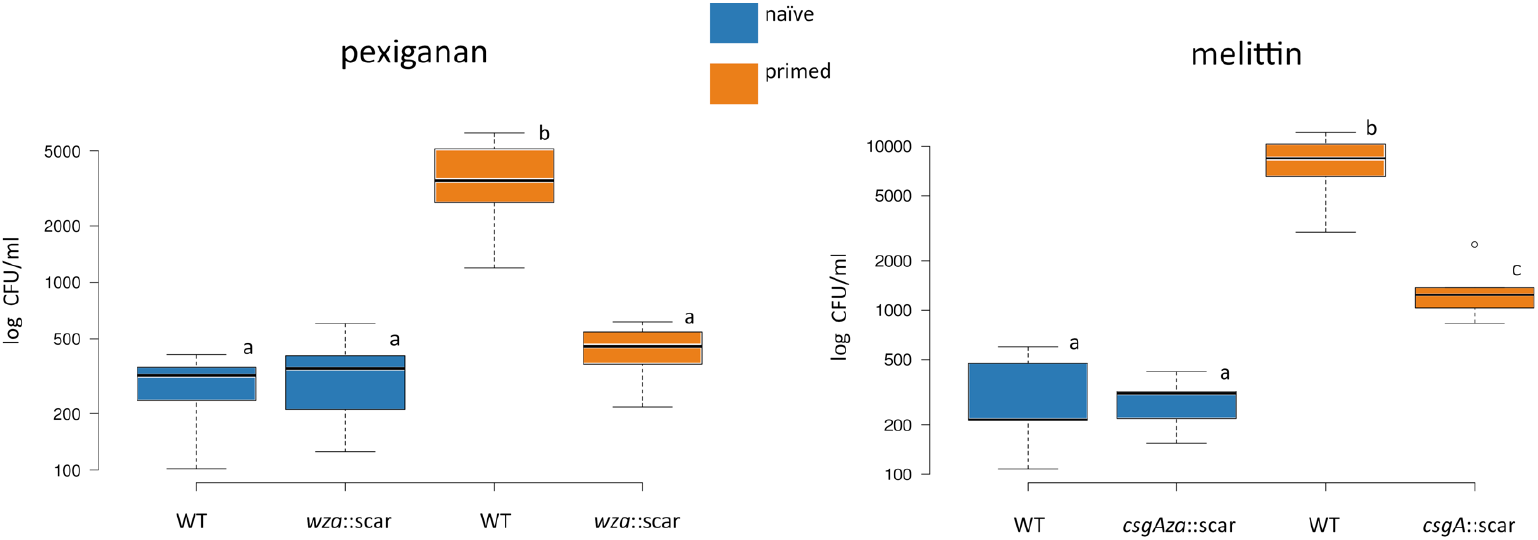
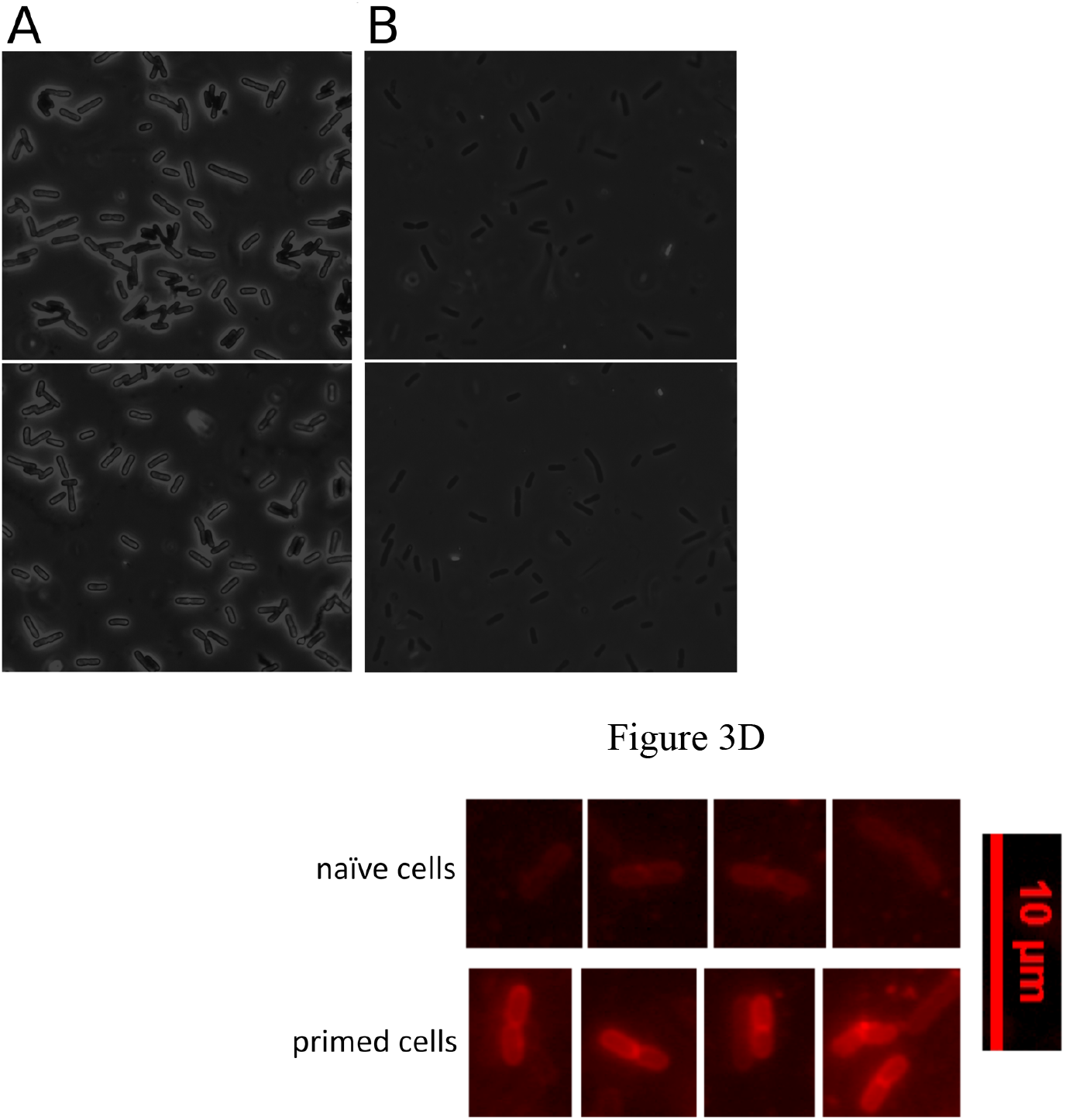
Physiological response to AMP priming. (A) SEM of E. coli MG1655 treated with 1/10xMIC (priming concentration) of pexiganan (top row) and melittin (bottom row) and non-treated bacteria (middle row, control). No apparent difference was noticed between melittin-treated cells and control. In the case of pexiganan, the treated cells tend to aggregate, a phenotype that consistent with the presence of colanic acid. Red arrows represent potential areas altered by the capsule of colanic acid that collapse under drying processing necessary for the preparation. Bacteria were observed with different magnifications ranging from 3000× to 40000X.(B). Bacterial mutants in *csgA* (curli mutant) and *wza* (colanic acid mutant) were exposed to the AMPs melittin and pexiganan respectively (10xMIC) after priming. Colony forming units (CFU) were determined after two hours of exposure. Boxplot show data tested by repeated-measures of one-way ANOVA and Dunnetts’ tests. For each case, equal letter represents no statistical differences while differing letter detonates significant ones (for p<0.05). (C). *E. coli* MG1655 treated with 1/10xMIC (priming concentration) of pexiganan (left) and non-treated bacteria (right, control) observed under phase contrast optical microscopy. The specimens consisted of cells suspended in a 0.1 % solution of nigrosin to create a strong contract to visualize the colanic acid capsules. Bacteria were observed with magnifications 1000× after exposure to a trigger stimulus. (D). Detection of curli production by primed cells after exposure to a killing dose of melittin (10xMIC). Curli production is only detected in small proportion of primed cells (survival fraction). After treatment, we exposed the cells to ECtracer™680 (Ebba Biotech, Sweden), a red fluorescent tracer molecule for staining of curli.

After lethal exposure to melittin and pexiganan we observed a high degree of heterogeneity regarding the killing of individual cells in primed bacteria, with many cells surviving the killing concentrations (Fig. 4 A). By contrast, almost all the control bacteria were killed (Fig. 4 A). As mellitin and pexiganan both damage the membrane, which ultimately leads to cell death, the live-dead stain despite it’s limitations, seems to be highly appropriate here. Primed cells also seem to aggregate, with a stronger effect in pexiganan-treated cells compared to melittin-treated cells (Fig. 4 A).

**Figure 4.**
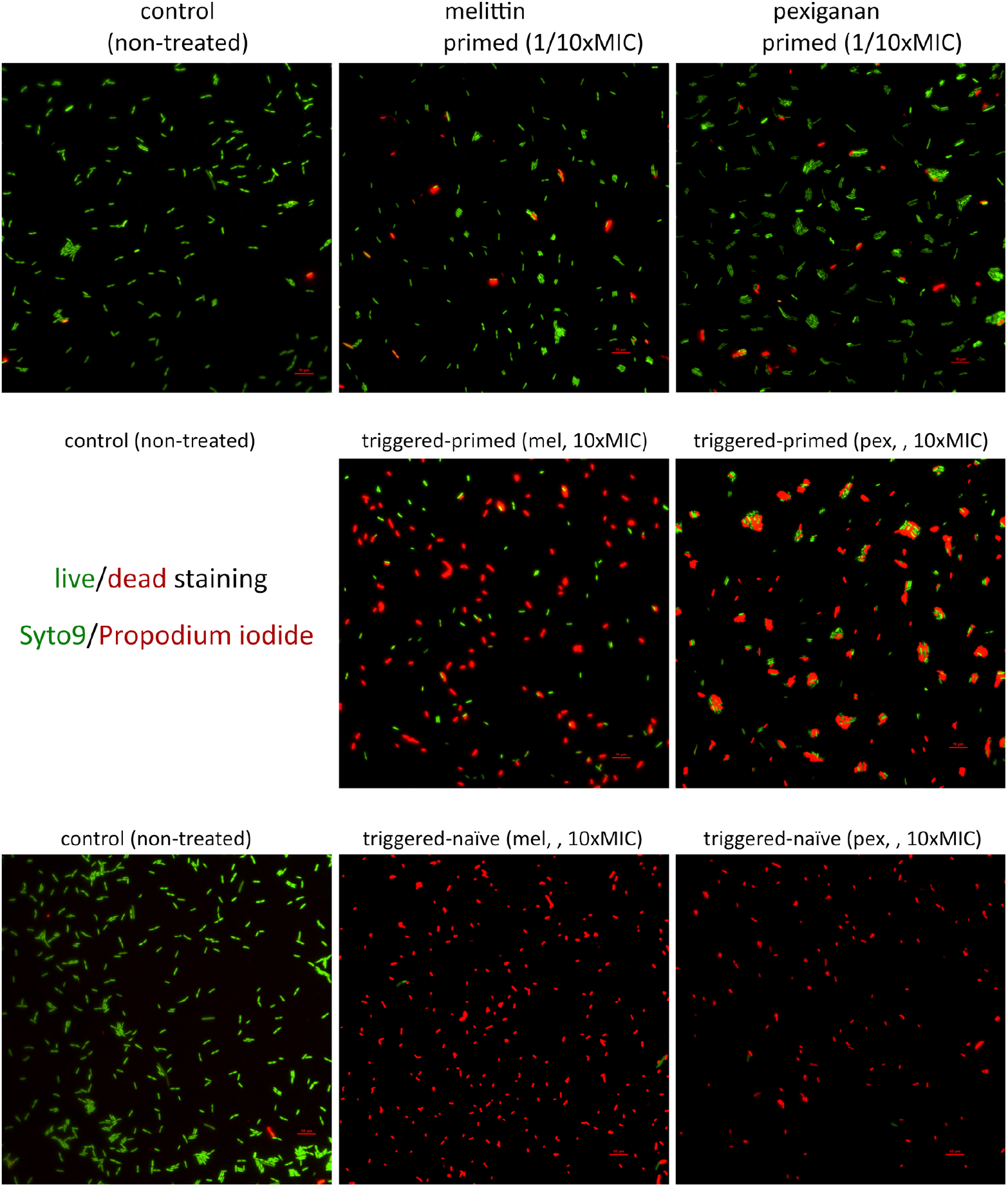
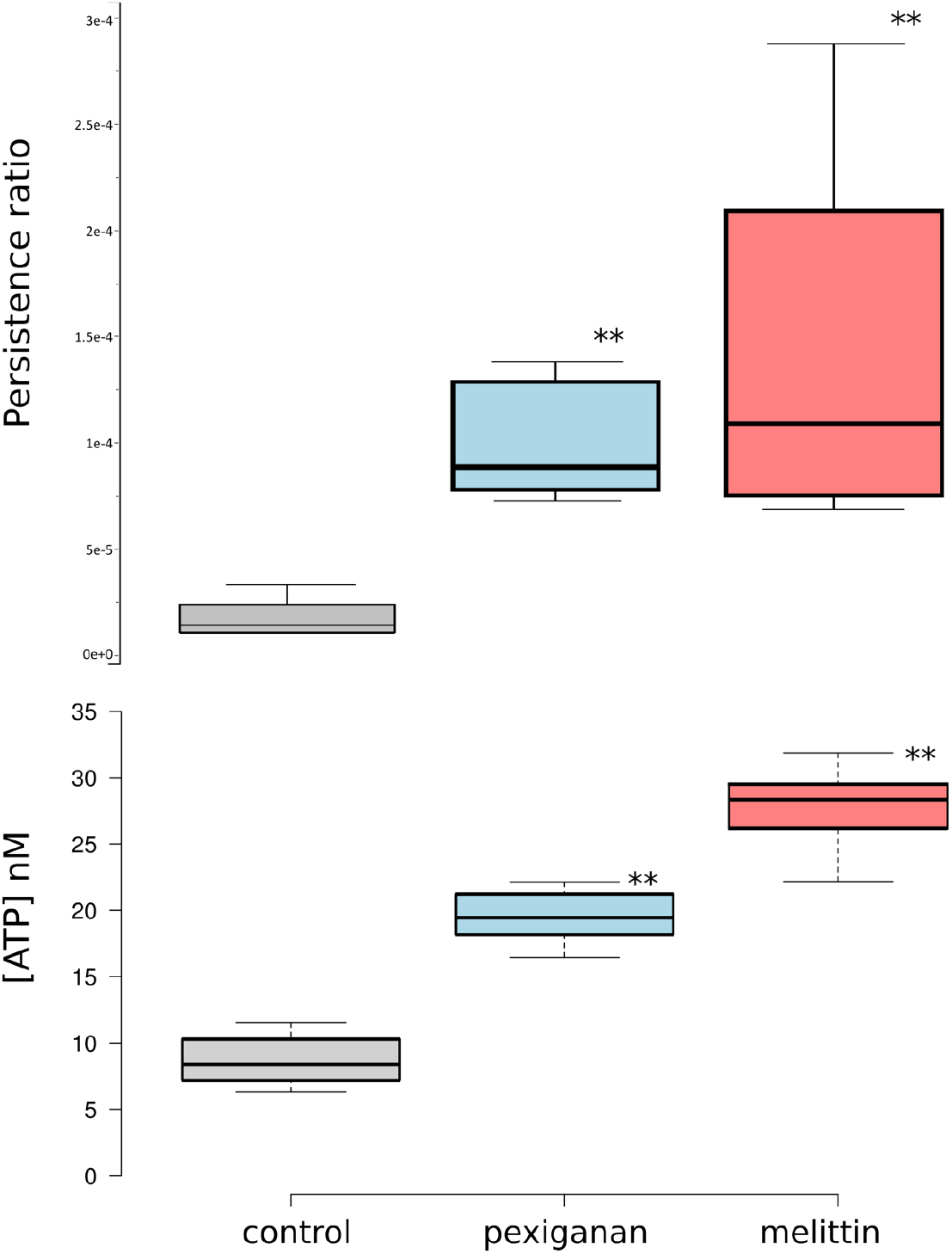
Cell viability and persister formation after AMP priming. (A) Cell viability after treating with priming and trigger doses of melittin and pexiganan were determined using the live/dead BacLight Bacterial Viability Kit (Thermo Scientific, Germany) on chip as described in M&M. After priming during 30 minutes, the treatment was removed by perfusing fresh MHB. The fluorescence signal was analyzed via a using excitation at 485 nm and emission at 530 nm for green fluorescence (Syto9) and using excitation at 485 nm and emission at 630 nm for red fluorescence (propidium iodide). (B) The priming concentrations of pexiganan and melittin, respectively, increased the persister number as determined by treated primed and naïve populations (10^8^ CFU/ml) with ciprofloxacin (10× MIC, 2μg/ml). This increase in persistence co-occurs with ATP leakage (determined in the supernatant of the culture medium).

To determine if priming stimulates persistence we used an assay based on the observation that a decrease of intracellular ATP increases persistence under the antibiotic ciprofloxacin (*32*). Because ATP leakage is a hallmark of AMP-treated bacteria (*33*), we exposed melittin and pexiganan primed populations and controls to ciprofloxacin (*34*). This resulted in a highly significant increase in the number of persisters (Fig. 4 B). The level of leaked ATP in the culture supernatant was significantly higher in primed bacteria for both AMPs as compared to controls (Fig. 4 B). The pre-treatment with melittin or pexiganan does not change the MIC of *E. coli* to melittin, pexiganan or ciprofloxacin consistent with the definition of persisters (*17*).

### A mathematical model predicts resistance evolution of primed cells

To quantify the influence of priming on bacterial tolerance and persistence, phenomena inherently linked to the growth dynamics and subpopulation structure, population dynamical models that capture tolerance and persistence are required. We build on a two-state population model previously developed (*25*) to describe bacterial antibiotic persistence (Fig. 5 A, B). This model assumes that bacteria exist in two phenotypic states, normal cells (*N*) and persisters (*P*). The two subpopulations *N* and *P* differ in their susceptibility to AMPs, a difference that is implemented as differing net growth rates for the given amount of AMPs (A), with *r_N_(A)* and *r_P_(A)*, respectively. Bacteria switch from subpopulation *N* to *P* with the rate *s_N_* and back with the rate *s_P_*.

**Figure 5:**
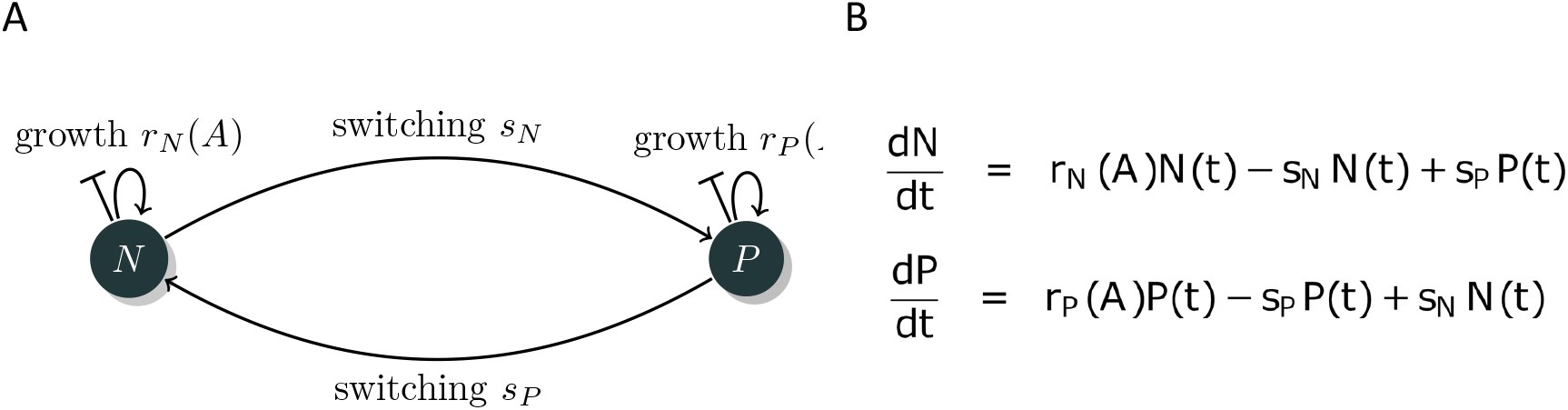
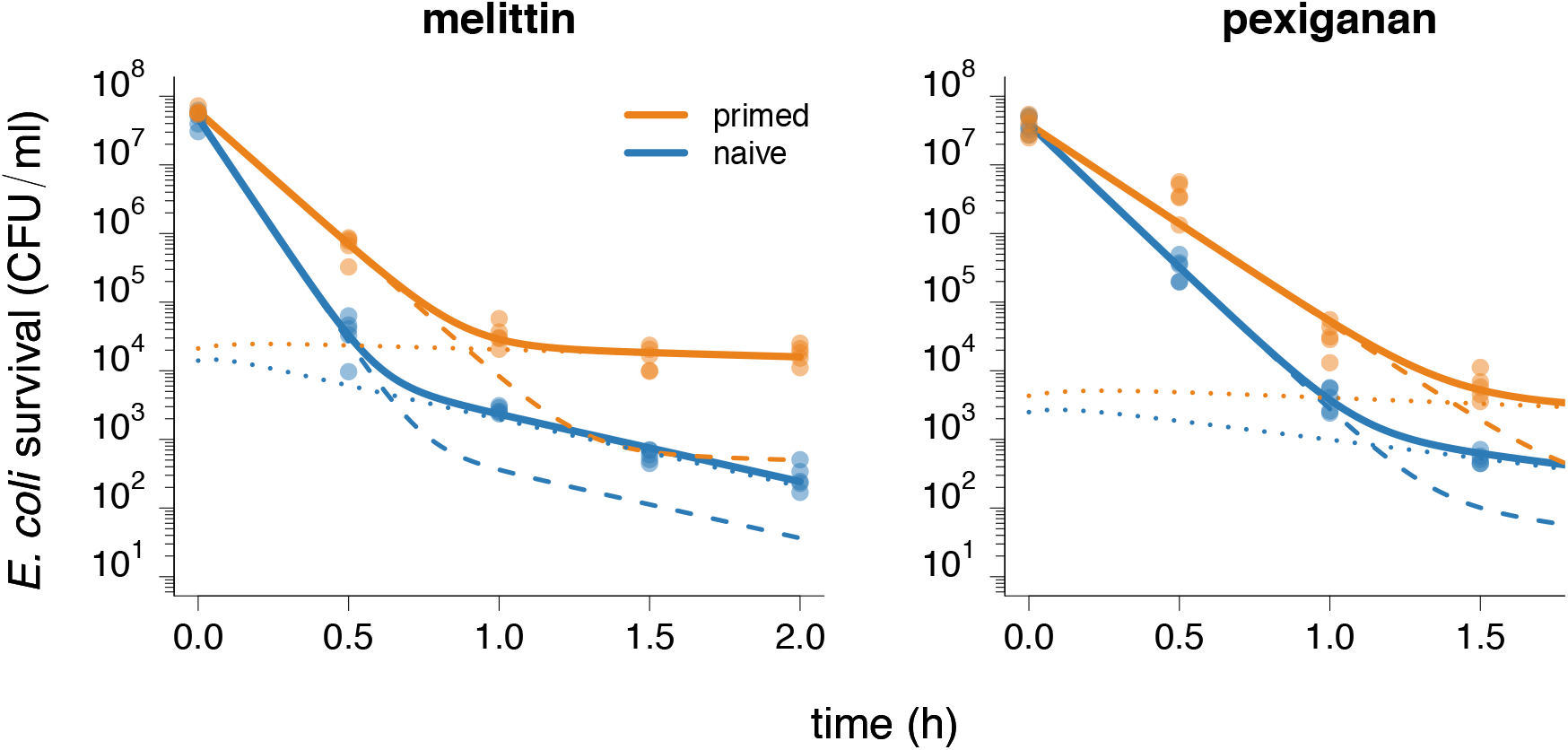
The two-state model describes time-kill data. (A) The two-state model (adapted from *17*) consists of two subpopulations, normal cells (N) and persister cells (P) and is parameterized with the growth rates r_N_(A) and r_P_(A), which are depended on the concentration of AMPs (A) in the system, and the switching rates sN, and sP. Each subpopulation is described with an ordinary differential equation (B), each of which describes the change of the respective subpopulation over time. For each antimicrobial, melittin (C) and pexiganan (D), we fitted the model to the data of naive (blue) and primed (orange) bacterial populations (see also Fig. 1) individually. The continuous lines represent the total bacterial population B(t), with B(t) = N(t) + P(t), and the dashed and dotted lines represent the subpopulations N(t) and P(t), respectively. Bacteria primed with melittin have an increased net growth rate r_N_ and decreased s_P_ compared to the naive populations. In the case of pexiganan, the parameter r_N_ is significantly higher in primed compared to naive populations. For an overview of the fitted parameters, see table S 5.

We quantified the model parameters by fitting the analytical solution of the model equations (Fig. 5 B) to the time-kill datasets of melittin and pexiganan. In a preliminary analysis with fitting all four parameters (rN, sN, rP, and sP), the net replication rate of persister cells, rp, was not significantly different from 0 (Fig. S 4). We therefore fixed rP to 0. The resulting fit described our data well (Fig. 5 C, D, and Table S 5). We found that priming affected two of the three estimated model parameters (Fig. S 4): rN increased, which translates into increase in tolerance, and sP decreased. Together, the increase in rN and decrease in sP result in higher persistence levels (Fig. 5 C, D).To assess the influence of priming on possible treatment success and resistance evolution, we extended a previously developed modeling framework to predict resistance evolution (*9*) by a persistent subpopulation (Fig. S 5). We then explored the effects of priming on tolerance and persistence, individually and in combination, which would be challenging empirically. Using our predictive framework, we investigated the effect of priming on survival based on a zero-order pharmacokinetic profile (*9*, *26*) (Fig. S 6) and with parameterized pharmacodynamics functions (Fig. S 7).

We found that survival of the population was highly dependent on tolerance (Fig. 6 A, B, S 8). The presence of persistent cells alone only marginally increased time until clearance. When we implemented the observed decrease in switching rate s_P_, the time until clearance of the bacterial population is extended at high treatment intensities (high concentrations A_max_). The results do not qualitatively change for larger pharmacokinetic decay rates (k), typical for AMPs (Fig. S 9). Taken together, an increase in tolerance alone resulted in higher survival independent of persister cells. An increase in persister cells further increased survival at high antimicrobial concentrations.

**Figure 6:**
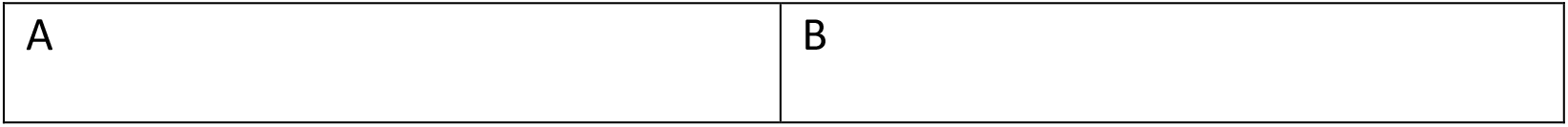
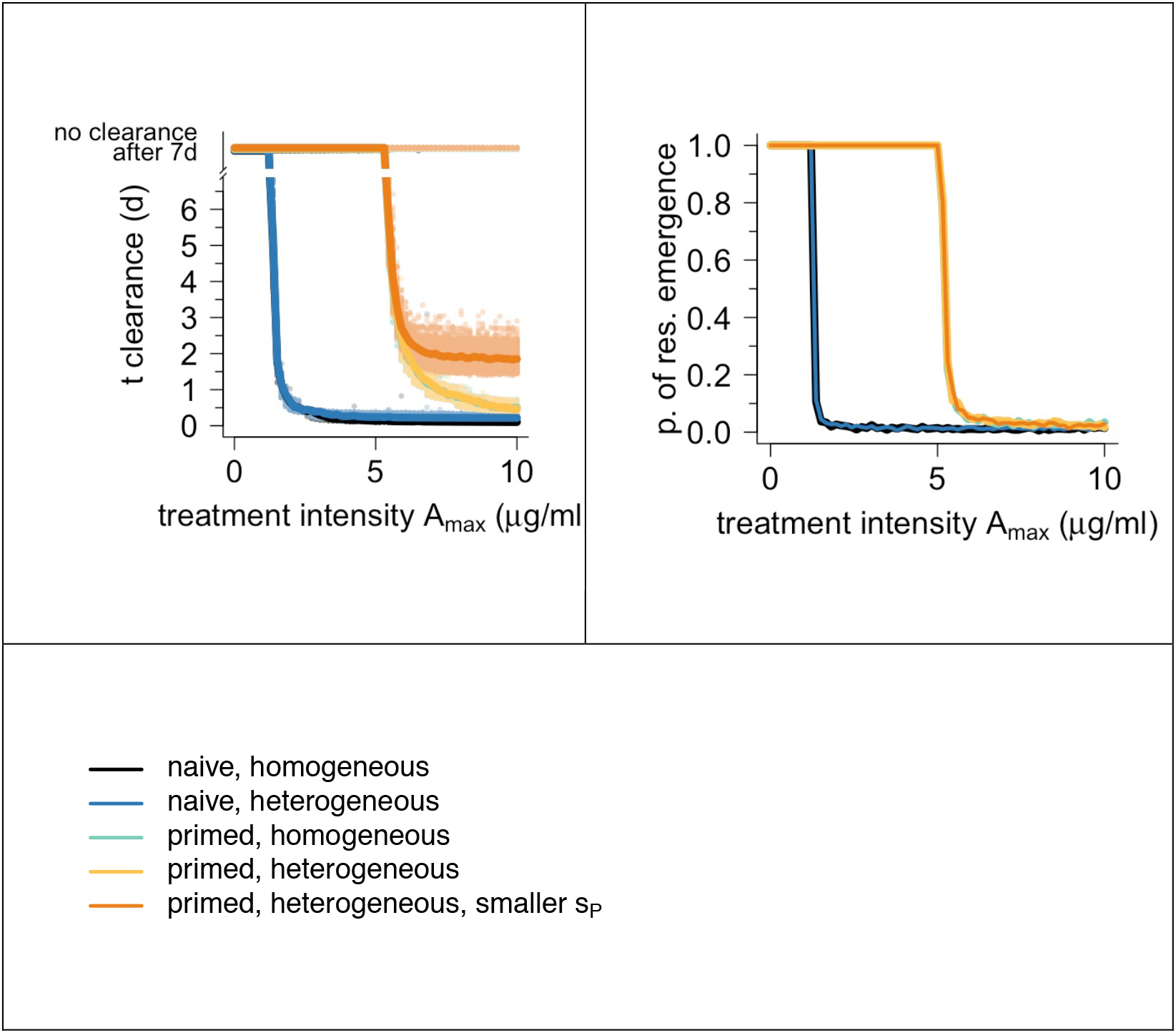
Influence of priming on time until clearance and resistance evolution. We extended a previously developed PKPD framework (*9*) to include persistence (Fig. S 5). With this framework, we estimated PD curves (Fig. S 7) for primed and naïve bacteria and the estimated switching rates (Fig. 5) to predict (A) time until clearance and (B) probability of resistance evolution. We simulated primed bacteria (primed with heterogeneous population consisting of N and P subpopulations, indicated as “primed, heterogeneous, smaller s_P_”) and naïve bacteria (naïve, heterogeneous). In addition, we simulated dynamics without persistence for both primed and naïve bacteria (“primed homogeneous” and “naïve homogeneous”) and dynamics of primed populations with switching rates of the naïve population (“primed, heterogeneous”). In (A), each dot is an individual run. In both (A) and (B), the line denotes the average of all runs. No clearance means that simulated treatment could not reduce bacteria population < 1 cell within 7 days of treatment. Here, we only show the results of melittin (results of pexiganan are qualitatively highly similar, see Fig. S 8). All parameter values used in the mathematical model are listed in table S 5.

Next, we assessed how priming affects resistance emergence. Generally, resistance evolution depends on the population size, mutation rate and the replication rate. While priming increases bacterial survival, hence the population size, it also increases the number of persisters that do not replicate and hence are a poor source of resistant mutations (Fig 1, Fig. S 10). Our simulations revealed that the beneficial effects of priming on survival due to increased persistence did not translate into an increased probability of resistance emergence and establishment (Fig. 6 B, Fig. S 8, 9). The probability of resistance emergence was mainly influenced by the effect of priming on tolerance.

## DISCUSSION

We find that sublethal dosing of the AMPs melittin and pexiganan primes bacterial cells to increase both tolerance and persistence. As sublethal concentrations of antimicrobials (AMPs and antibiotics) are common in the environment and in hosts/patients, a priming response, if it is a widespread phenomenon, would have significant consequences. Antimicrobials that prime inevitably would induce the formation of tolerant and persisting cells. This should apply to situations of induced immune responses, but equally to drug treatments. In short, generating or increasing populations of phenotypic resistant bacteria, would be inevitable in many situations, and would represent a serious obstacle to clearing bacterial infections.

The priming response we report is mediated by rather general physiological resistance mechanisms. The molecular basis of the induction of AMP tolerance and persistence is based on modifications of bacterial envelopes involving either curli production under melittin treatment, or colanic acid production after pexiganan exposure. Both these mechanisms protect the cells by shielding them from AMPs, either making the access to the outer membrane more difficult and/or capturing AMPs, giving scope for cross-resistance. These physiological resistance mechanisms are not antimicrobial specific and hence likely to be also induced by other antimicrobials.

Interestingly, the activation of both pathways shows different dynamics in biofilm formation **(*35*)**. Curli and colanic acid are important components of the biofilm matrix, curli for the primary adhesion and colanic acid for the biofilm structure **(*35*)**. Triggering their expression by sublethal levels of AMPs, could potentially catalyse biofilm formation. Within a host, if the immune system fails to clear the pathogens, the AMP-priming effect may thereby favour the transition from acute to chronic bacterial infections, where biofilms prevail.

In natural systems of host-microbe interactions, priming plays a role in facilitating colonization of AMP rich environments **(*11*)**. Priming by AMPs also plays a role in infection vectors: in the flea gut the PhoQ-PhoP system is induced in *Yersinia pestis*, the infective agent causing plague, by AMPs leading to biofilm formation that enhances transmission to the final host **(*36*)**. It is not clear as yet if phenotypic AMP-resistance will facilitate opportunistic infections in a way similar to genetic AMP-resistance, as has been shown for genetic AMP resistance in *S. aureus* **(*37*)**, but in the light of our results it seems likely.

Toleranc and persistence can drive the evolution of genetic resistance **(*10*)** *(18).* We find that, while priming *E. coli* with AMPs results in increased tolerance and persistence, the main driver of resistance evolution in our model is increased tolerance. Resistance evolution is a product of mutation supply and strength of selection. The selection in our scenarios is strong, but mutations are only supplied from the subpopulation of tolerant cells, not from the persister cells, as they are metabolically inactive. So, while the persisters do not contribute to resistance evolution, they do provide a source for secondary infection, once the immune system overexpression ceases or alternatively, when the antimicrobial drug is removed. In antibiotic resistance evolution, by contrast, persisters have shown to contribute to resistance evolution **(*38*)**. Part of this seems to be explained by increased mutagenesis caused by the antibiotics **(*38*)**, an aspect we explicitly did not model as AMPs do not increase mutagenesis **(*39*)**.

Our combined theoretical and empirical results suggest that in hosts the optimal strategy of AMP-deployment would be a very fast increase in concentration to avoid priming and the subsequent development of phenotypic resistance through both, increased persistence and increased tolerance. Such fast increases of AMPs in specific locations are realized in some important natural situations. During insect metamorphosis, when the gut is renewed in the pupa, a cocktail of AMPs and lysozymes is discharged into the gut **(*40*)** and references therein), resulting in a quick reduction of bacterial numbers by orders of magnitude. When infections persist, one possible solution is a long-lasting immune response **(*41*)** that deals with a recurrent infection that could potentially be caused by persisting cells. These natural situations also have parallels in the medical application of antimicrobials. We propose that a fast increase of antimicrobial concentration, as for example in the intra-venous application of antibiotics, should also reduce the probability of persister formation via priming.

Long-lasting antimicrobial exposures are prevalent in natural systems. This is at odds with the observation that long-lasting drug treatments, at least in the case of single drug applications, select for drug resistance. Therefore, at first glance extended treatments do not seem to be a good strategy to manage persisters. In the natural context of AMPs, as they are always expressed as cocktails, this is very likely not an issue and would suggest the use of combination therapies.

## Acknowledgments

We are grateful to Sophie Armitage, Charlotte Rafaluk-Mohr and Olivia Judson for comments on the manuscript.

## METHODS

### Bacteria and growth conditions

The *E. coli* MG1655 was used as bacterial models for all experiments. All cultures related to antimicrobial tests were carried out in Mueller-Hinton I Broth (Sigma). For genetic manipulations (*E. coli* BW25113). All the strains and their derivatives were routinely cultured in Lysogeny Broth (LB medium) or SOB, supplemented with antibiotics when appropriate. Other constructed and strains used in this study are listed in table S 6 of this section.

### Minimal inhibitory concentration (MIC)

MICs were determined according to CLSI recommendations by a microdilution method in MHB with some modifications. Inoculum size that was adjusted to 2×10^7^ CFU/ml from a regrowth of overnight cultures to be consistent with the downstream experiments. The MIC was defined as the antimicrobial concentration that inhibited growth after 24 h of incubation in liquid MH medium at 37°C. Polypropylene non-binding plates (Th. Geyer, Germany) were used for all experiments. The MIC was considered as the antimicrobial concentration that inhibited growth after 24 hours of incubation in liquid medium at 37°C.

MIC was also determined similarly as above for primed bacteria after two and a half hours of exposure to priming concentrations. The values were compared with naive controls for both antimicrobial peptides, melittin and pexiganan, and for ciprofloxacin.

### Priming experiments

Starting from 1×10^8^ CFU/ml in mid exponential growth, where bacteria were exposed (stimulus) to 1/10 MIC of melittin or pexiganan during 30 minutes at 37°C with soft shaking. The tubes were centrifuged at 4000 × g for 10 minutes and allowed to recover for 60 minutes. The cells were challenged (trigger of priming response) with a concentration equivalent to an 10× MIC. The cultures were diluted and plated to determine cell viability. Five biological replicates were generated. Non-treated cells were used as a control and also harvested during mid exponential growth.

### Activity of melittin and pexiganan from the supernatant of challenged cells

Similarly, to the priming experiments, 1×10^8^ CFU/ml of bacteria were exposed 10× MIC (supernatant I) of melittin or pexiganan during 150 minutes at 37°C with soft shaking. The tubes were centrifuged at 4000 × g for 10 minutes, the supernatant were collected and centrifuged again for 20 000 × g for 30 minutes. The new supernatant was filtrated using 0.22 μm sterile filters (Sigma Aldrich, Germany) and used immediately (supernatant II). In parallel, exponentially growing cultures containing 1×10^8^ CFU/ml per tube were centrifuged at 4000 × g for 10 minutes, the supernatants were removed by sterile aspiration and the pellets were re-suspended in equal volumes of the supernatant II and incubated during 150 minutes at 37°C with soft shaking. Samples from each tube were from both, supernatant I and supernatant II, taken every 30 minutes and diluted and plated to determine cell viability. Five replicas per culture were used. Data points from time-kill experiments from bacteria treated with the supernatant I (first round) and the supernatant II (second round) were compared to determine changes in the activity of the AMPs either by degradation or adsorption.

### Persister antibiotic survival assay

Bacterial cultures were inoculated at 1:100 from a 16-hour overnight culture into MHB medium. Cells were grown for 2 h to reach approximately 2 × 10^8^ CFU/ml). The cultures were treated with priming concentrations (1/10 MIC) of melittin and pexiganan for 30 minutes. Non-treated cultures were used as control. All cultures were washed and centrifuged twice to remove the treatment. The supernatants were used to determine ATP concentration using a Molecular Probes ATP Determination Kit (Thermo Fisher Scientific, Germany). Bacteria were resuspended in equal volume of fresh medium and an aliquot from each culture was taken to determine the number of bacteria at t=0 by diluting and plating in MHB agar. Following, ciprofloxacin was added for a final concentration of 2 μg/ml to treated tubes and to non-treated AMPs control. The cultures were incubated for four hours. Then, bacteria were washed twice with 0.9 % NaCl and plated on MHB agar to determine the counts of survival fractions. The percent survival was calculated as the ratio of cells before and after the treatment as described previously (*34*). Briefly final CFU/CFU at 0 h) × 100. The results are presented as the average from 5 independent replicates.

### Construction and verification of deletion mutants

We inactivated the major curli subunit protein gene *csgA* and the colanic acid precursor gene *wza*. Although both pathways involve many genes, the removal of these two components impair the production of both substances respectively. These mutants were generated in *E. coli* K‐12 strain MG1655 following the methodology described elsewhere (*42*) with some modifications because we used as template the genomic DNA of the Keio collection (*43*). Briefly, we extracted genomic DNA from the mutants *csgA::Kn* and *wza::Kn* of the Keio collection (*E. coli* BW25113) and amplified by PRC the flanking regions of the kanamycin resistance cassette disrupting both genes and including an appropriate homology sequence. For the *csgA* mutant we used the primers 5’-GATGCCAGTATTTCGCAAGGTG-3’ and 5’-GGTTATCTGACTGGAAAGTGCC-3’ while primers 5’-TAGCGTGTCTGGATGCCTG-3’ and 5’-CCACTTTCAGCTCCGGGT-3’ were used for *wza*. The PCR products were purified and electroporated in the *E. coli* MG1655 carrying a red recombinase helper plasmid, pKD46. The strain was grown in 10 ml SOB medium with ampicillin (100 μg/ml) and L-arabinose at 30°C to an OD_600_ of ~0.5 and then made electrocompetent by washing and centrifuge them with a cold solution of glycerol 10%. Competent cells in 80 μl aliquots were electroporated with 200 ng of PCR product. Cells were added immediately 0.9 ml of SOC, incubated during 2 h at 37°C, and then 100 μl aliquots spread onto LB agar with kanamycin (30 μg/ml). The correct inactivation of genes was verified by PCR. The antibiotic resistant cassette (Kn) was removed for both mutants using the flippase plasmid pCP20.

### Transcriptome sequencing

The transcriptome sequencing of primed cells was determined on samples treated identically as described for the priming experiments. Total RNA from 10^8^ cell per sample was isolated using the RNAeasy Isolation kit (Qiagen, Germany). Traces of genomic DNA were removed from 10 μg of RNA by digestion in a total volume of 500 μl containing 20 units of TURBO DNase, incubated for 30 minutes at 37°C, immediately followed by RNeasy (Qiagen) clean-up and elution in 30 μl of RNase-free water. Following DNase treatment, RNA integrity was assessed using Agilent RNA 6000 Nano kit and 2100 Bioanalyzer instrument (both from Agilent Technologies). Total RNA was depleted from ribosomal RNA using the Ribo-Zero Depletion Kit for Gram-negative bacteria (Illumina, USA). Libraries were prepared using a TruSeq Stranded Total RNA library preparation kit (Illumina, USA) and were sequenced on a MiSeq platform. Transcript abundances were derived from pseudo-alignment of reads to the cDNA sequences from the ASM584v2 assembly of *Escherichia coli* MG1655 (ENA accession GCA_000005845.2) using Salmon version 0.7.2 with default parameters (*44*). Differential gene expression was analyzed using the R package DESeq2 (*45*) in conjunction with tximport (*46*). Pairwise contrasts were performed between the control and each AMP treatment with empirical bayesian shrinkage of both dispersion parameters and fold-change estimation. We defined genes as being significantly differentially expressed when the absolute fold-change in expression was greater than 2, at an FDR-adjusted p-value of less than 0.05. The variance-stabilizing transformation was used to remove the dependence of the variance on the mean and to transform data to the log2 scale prior to ordination using principal component analysis. Quality of RNAseq data were contrasted by Euclidian distance and symmetry of data reads distribution (Fig. S 11).

### Observation at single cell level

To observe cell reaction at single cell level during priming experiments, we used an ad hoc microfluidic device developed for this project. It consisted in a main channel for bacterial inoculation and medium perfusion and with several lateral compartments which dimensions are around 1.5 μm height (ensuring all bacteria are kept in focus) and square 200 μm width corresponding to a field size of the used microscope at 1000× magnification (Fig. S 12). The chip was designed in Autocad. We started the replication of our microfluidic chips from a custom made (Sigatec SA) silicon (SiO) master. This silicon master itself was first replicated in Smooth-Cast 310 (Bentley advanced material). Soft lithography produced the chips in PDMS (Sylgard Silicone Elastomer Base and Curing Agent mixed in 10:1 ratio). The PDMS chips were cured overnight at 75°C in an incubator. We punched an inlet and outlet hole for the laminar flow in each chip using a biopsy puncher of 0.5 mm (outer diameter). The chips were bonded to a glass cover slide (24×60 mm) after a 30-second air plasma treatment (PDC-002, Harrick Plasma). Before use, the assembled chip was treated for 15 seconds in air plasma and immediately injected it with filtered MHB medium for passivation. We left the activated chip to incubate for a least 1 hour before loading the bacteria. The devices were loaded to full capacity with a bacterial suspension containing nearly 2×10^8^ CFU/ml (exponentially growing bacteria, 0.5 OD_600_). Cell suspension was injected into the main channel of the chip using a bluntend 23G needle attached to a 1ml syringe. We centrifuged the loaded chip at 200 × g for 10 min using in-house adapters, checked the loading.

After we loaded the bacteria cells in the blind ending side channels, we connected the chip to a syringe pump (AL-6000, WPI, Germany) and placed the chip under an inverted microscope. A continuous laminar flow (100μl/h) of MHB through the central channel was maintained throughout the experiment (Fig. S 12). For the life cell imaging, after infusion with priming or triggering concentration of AMPs, we injected MHB supplemented with bacterial Live/Dead stain kit solutions (Thermo Scientific, Germany) for a final concentration of 0.1 μl/ml of MHB. We took pictures of at least 20 fields per treatment from independent side channels. Fluorescent images were taken of each field of view with simultaneous acquisition in red and green fluorescent channels during a time interval of no more than 2 minutes per treatment with a Nikon Ti-2 inverted microscope (Nikon, Japan). Cells were observed with the 100× objective and controlled by Nis Element AR software. The chip holder is temperature controlled at 37°C.

### Determination of melittin-induced curli

The production of curli due to melittin treatment was determined by using the fluorescent dye Ectracer™^680^ (Ebba Biotech, Sweden) that stain extracellular curli. Ectracer™^680^ was used according to the instructions of manufacturers. Bacterial cultures were treated on chip with priming and triggering concentrations of melittin as described for single cell observation section omitting the live/dead staining. After priming and triggering, the channels were perfused MHB supplemented with Ectracer™^680^ in a proportion of 1/1000 related to the medium. Cells were observed with the red channel fluorescence for Cy5 dye using a Nikon Eclipse Ti2 inverted optical microscope using the 100× oil objective. Two independent samples were prepared for each group (primed and naïve cells).

### SEM of *E. coli* treated antimicrobial peptides

Approximately 2×10^7^ CFU/ml *E. coli* MG1655 were treated with 1/10 MIC of pexiganan and melittin during 30 minutes. The cultures were concentrated 10 times by a quick centrifugation step of 1 minute at 8 000 g and resuspended in 1/10 of its own supernatant. and resuspension 10 μl drops were placed on a circular glass cover slip (1.5 cm of diameter). The drops were fixed with osmium tetroxide vapors during one minute and allow to dry in a laminar flow cabinet. The cover glasses were mounted on aluminum stubs using double-sided adhesive tape and coated with gold in a sputter coater (SCD-040; Balzers, Union, Liechtenstein). The specimens were examined with a FEI Quanta 200 SEM (FEI Co., Hillsboro, OR) operating at an accelerating voltage of 15 kV under high vacuum mode at different magnifications. At least 5 fields from two independent replicas.

### Statistical analysis

To analyze the priming data, we first tested if the dynamics of the depicted in the time-kill curves are biphasic.

We fitted the function

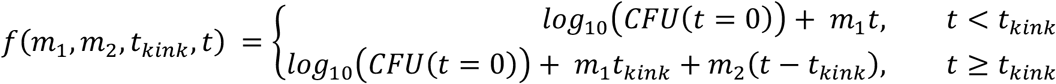

to the time-kill data of each AMP and for primed and naïve populations individually using a least square algorithm. Here, *t_kink_* is the time point, at which the population dynamics switch from the first phase to the second phase and *m_1_* and *m_2_* are the slopes of the first and the second decline, respectively. Note that *m_1_* is a direct measure of tolerance. The standard error (SE) was calculated as

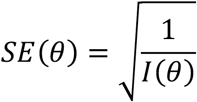

The parameter *θ* denotes to the estimated parameter values of *m_1_*, *m_2_*, and *t_kink_* and *I*(*θ*) is the expected Fisher information. The 95% confidence interval was calculated as *θ* ± 1.96 ∗ SE(*θ*).

We used an ANOVA by means of contrasts to assess if priming changes tolerance and persistence. For this, we tested if the decrease of the first phase (all data points with t < t_kink_) differed between primed and naive populations (test for differences in tolerance) and if the population size in the second phase (all data points with t > t_kink_) differed between primed and naive populations (test for differences in persistence). Both tests showed significant differences between primed and naive populations for both melittin and pexiganan (significance level: p < 0.05). We corrected for multiple testing with the Bonferroni-method.

### Population models

To describe bacterial population dynamics, we used the two-state model by Balaban *et al.* (*25*) (Fig. 5 A, B):

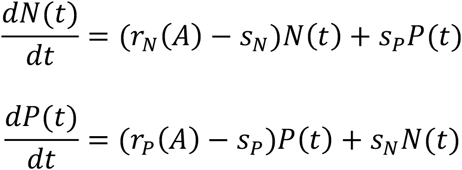

In this model, the bacterial population consists of two subpopulations, one with a normal phenotype, *N(t)*, and a second with a persister phenotype,*P(t)*. We denote the total bacterial population size by *B(t)*,with *B(t)* = *N(t)*+*P(t)*. The rate of change of the population is determined by the net growth rate of *N* and *P*, *r_N_* and *r_P_*, and the switching rate from *N* to *P*, *s_N_*, and the switching rate from *P* to *N*, *s_P_*. The parameter estimates of the net growth rates are dependent on the AMP concentration A, i.e. *r_N_(A)* and *r_P_(A).* analytical solution of this ODE system (*25*, *47*) is

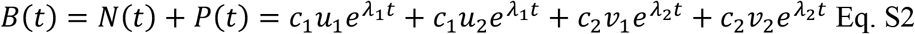

with

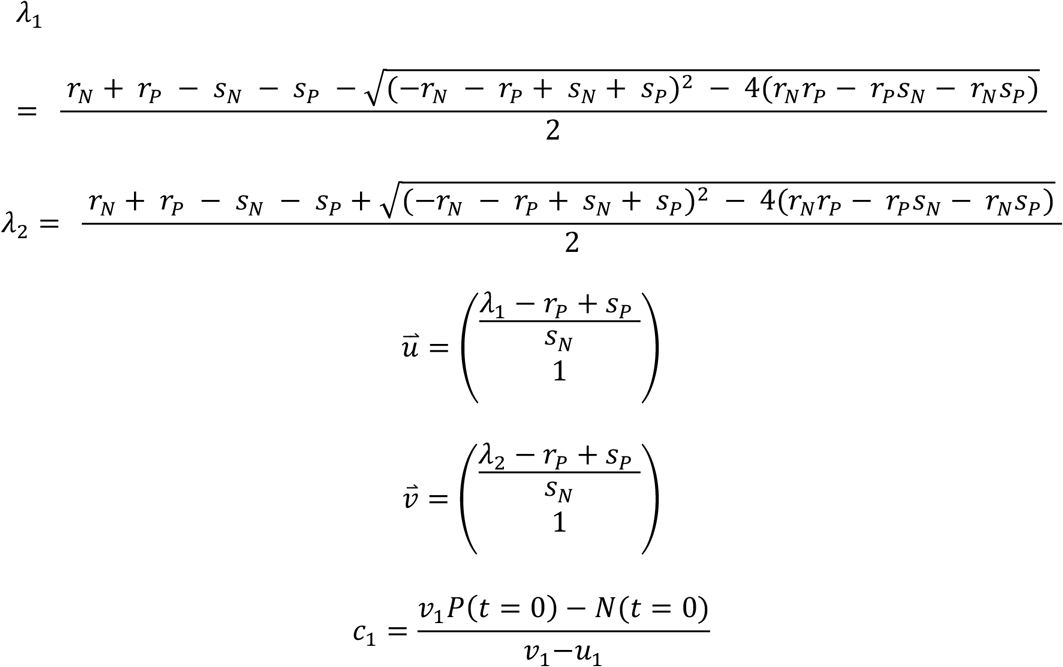

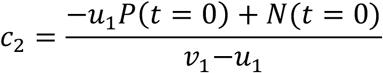

The model was fitted by a least square algorithm, minimizing the residual sum of squares of the data to the prediction. For the starting conditions (*N*(*t*=0), *P*(*t*=0)), we assumed that the ratio of *N*/*P* was at the equilibrium predicted by the model without antimicrobials when the exposure to lethal concentrations of AMPs started. *N(t=0)* and *P(t=0)* were therefore calculated using the eigenvector 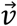 that corresponds to the largest eigenvalue of a system without antimicrobials. Here, we assumed that the parameter *r_N_* is equal the net growth rate in absence of antimicrobials, *r*_*N*_ = *ψ*_*max*_. The parameter *ψ*_*max*_ was estimated based on the time-kill curve of bacterial population that grow in absence of antimicrobials (see below and Fig. S 11). The eigenvector contains information about the ratio of N and P for 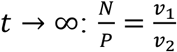. Resulting, 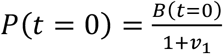 and *N*(*t* = 0) = *B*(*t* = 0) − *P*(*t* = 0). *B*(*t* = 0) was estimated from the data. Confidence intervals were calculated as described above. In a pre-analysis, we used 4 free model parameters that were fitted: *r_N_*, *r_P_*, *s_N_* and *s_P_*. The parameter *r_P_* was not significant from 0 (Fig. S 8). Therefore, we set the parameter *r_P_* to 0 and fitted the remaining 3 parameters to the data (Fig. S 9, table S 5).

### Tolerance and persistence in terms of model parameters

The measure of tolerance is the slope *m_1_*. Komarova and Wodarz (*47*) showed that the slope can directly be linked to the population model parameters. In our notation,

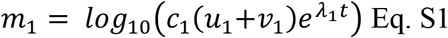

Note that the first phase is mainly influenced by *r_N_* (Fig. S9), therefore, *m*_1_ ≈ *r*_*N*_.

Persistent cell numbers at time t were calculated with the analytic solution:

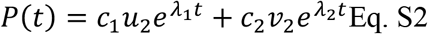

### Pharmacokinetic and pharmacodynamic function

We used the pharmacokinetic function *A*(*t*) = *A*_*max*_*e*^−*k*(*t−n*)^, with 8*h* ∗ *n* ≤ *t* ≤ 8*h* ∗ (*n* + 1) and *n* = 0,1,2 … described previously elsewhere (*26*). In our simulations, we fixed the decay parameter *k* and varied the drug input *A_max_*. To describe the effect of the AMPs on the bacterial population, we used the pharmacodynamic (PD) function *ψ*(*A*) (*26*, *48*), with

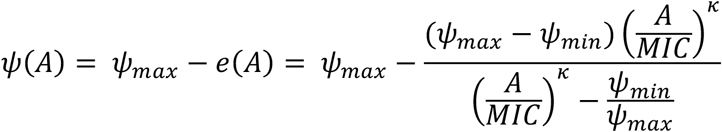

The parameter *ψ*_*max*_ describes the net growth rate in absence of antimicrobials (*ψ*_*max*_ = *ψ*(*A* = 0)). The antimicrobial effect *e*(*A*) is dependent on the antimicrobial concentration and is the defined with *ψ*_*max*_, *ψ*_*min*_, the net growth rate in presence of large amounts of antimicrobials (*ψ*(*A* → ∞)), with the MIC, the antimicrobial concentration that results in no growth (*ψ*(*A* = *MIC*) = 0) and with *κ*, which determines the steepness of the PD curve.

The PD function was fitted to the time-kill curves (Fig. S6), as described by Regoes *et al.* (*26*). In short, we used log-linear regressions of the time kill curves within the time-points 0h and 1h to estimate the change of the bacterial population over time, i.e. the slopes of the log-linear regression. We fixed the parameter *ψ*_*max*_ and fitted the 3 remaining parameters of the PD function with the *Markov-Chain-Monte-Carlo method*.

### Stochastic simulations

To simulate resistance evolution with stochastic simulations, we expanded a previously developed framework for bacterial population dynamics (*9*). The framework models bacterial population dynamics exposed to changing levels of antimicrobials and allows for resistance evolution. In the simulations, the change in population size of a sensitive strain *S*, with *S* = *N*+*P*, and of a resistant strain *R* were described with the following ODE system:

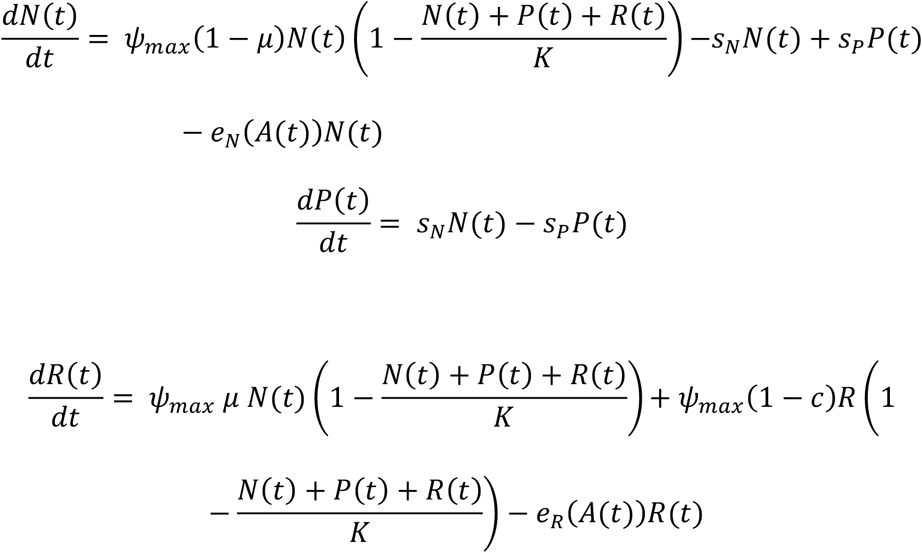

Here, the replication rate is assumed to be equal to the maximum net growth rate *ψ*_*max*_. The effect of the antimicrobial *e*_*N*_(*A*) is explained above. Note that we assume that bacteria in class P do not grow and are not affected by antimicrobials. We also assumed that the switching rates are constant. To describe the effect of an antimicrobial on the strain *R*, *e*_*R*_(*A*), we use the same parameter set than with *e*_*N*_(*A*), except for the *MIC*: 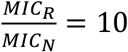. In the simulations, we differentiated between the following cases: (i) naïve bacteria, homogeneous population (no persistence), (ii) naïve bacteria, heterogeneous population, (iii) primed bacteria (increase in *r_N_*), homogeneous population, (iv) primed bacteria (increase in *r_N_*), heterogeneous population (v) primed bacteria (increase in *r_N_* and decrease in *s_P_*). The stochastic simulations were run 1000 times for each antimicrobial, for each case, and for a variety of input antimicrobial concentrations *A_max_*. All parameter values are listed in table S 6. The intensity is the input antimicrobial concentrations *A_max_*. The simulations were run for t=7d. Time until clearance and probability of resistance evolution were calculated as the mean of the value over the 1000 simulations.

### Implementation

Statistical testing, simulations and plots were done in R version 3.3.2 (*49*), using Rstudio version 1.0.143 (*50*). We used the following R-packages: (i) for plotting: sfsmisc (*51*), (ii) for fitting the PD function: rjags(iii) for stochastic simulations: adaptivetau (*52*). We used Mathematica version 11.0 (*53*) to determine the analytical solutions of the population models.

## Supplementary tables

**Table S 1.** Minimal inhibitory concentration of the two antimicrobial peptides melittin and pexiganan and the antibiotic ciprofloxacin used in this work. The MIC was determined to be used as a concentration reference but it was also determined after priming treatment to show that the observed enhanced survival is based on a phenotypic changes and not mutations.

| Antimicrobials | MIC (µg/ml) pre-priming | MIC (µg/ml) post-priming |
| --- | --- | --- |
| melittin | 2 | 2 |
| pexiganan | 1 | 1 |
| ciprofloxacin | 0.125 | 0.125 |

**Table S 2.** Slopes of fitting the biphasic function with the slopes *m_1_* and *m_2_* to the data on a log_10_ scale. Significant differences between slopes of primed and naïve population counts are indicated with asterisks for each antimicrobial.

| Antimicrobial | Treatment | Slope |  |  |  |
| --- | --- | --- | --- | --- | --- |
| | | $m_1 (\pm \text{SE})$ | | $m_2 (\pm \text{SE})$ | |
| melittin | Primed | -3.9 ($\pm 0.6$) | * | -0.3 ($\pm 0.4$) | |
| | Naïve | -6.3 ($\pm 0.6$) | | -1.0 ($\pm 0.4$) | |
| pexiganan | Primed | -2.9 ($\pm 0.3$) | * | -0.6 ($\pm 0.894$) | |
| | Naïve | -4.1 ($\pm 0.3$) | | -0.3 ($\pm 0.9$) | |

**Table S 3.** Priming response to pexiganan (a) and melittin (b) (0.1xMIC, 30-minute treatment) *See excel files*

**Table S 4.** Results of fitting the two-state model to bacterial population dynamics in the presence of melittin and pexiganan and the classic exponential population growth model to dynamics in the presence of ciprofloxacin and ampicillin (rounded values).

| Antimicrobial | Treatment | $r_N (\pm \text{SE})$ | $r_P^1$ | $s_N (\pm \text{SE})$ | $s_P (\pm \text{SE})$ |
| --- | --- | --- | --- | --- | --- |
| melittin | Primed | -8.96 (0.72) | 0 | 0.00085<br>(0.00098) | 0.27 (0.61) |
|  | Naïve | -15.06 (0.86) | 0 | 0.00128<br>(0.00095) | 2.25 (0.45) |
| pexiganan | Primed | -6.68 (0.32) | 0 | 0.00027<br>(0.00059) | 0.40 (1.10) |
|  | Naïve | -9.61 (0.40) | 0 | 0.00021<br>(0.00025) | 1.26 (0.66) |

<sup>1</sup> Parameter $r_P$ is fixed at 0 due to results of the preliminary analysis, in which the parameter was not significantly different from 0 (see Figure S 8).

**Table S 5.** Parameter values used as input for the stochastic simulations.

| Parameter | Description | AMP | Value |
| --- | --- | --- | --- |
| $k$ | AMP degradation rate | | 0.3 |
| $\mu$ | Mutation rate | | $1 \cdot 10^{-9}$ |
| $K$ | Carrying capacity | | $5 \cdot 10^9$ |
| $\psi_{max}$ | Maximal net growth rate in absence of antimicrobials | melittin | 2.13 |
|  |  | pexiganan | 2.06 |
| $\psi_{min}$ | Minimal net growth rate in presence of large amounts of antimicrobials | melittin | -8.86/ -9.89<sup>§</sup> |
|  |  | pexiganan | -36.87/-66.80<sup>§</sup> |
| $\kappa$ | Hill parameter | melittin | 1.3/1.1<sup>§</sup> |
|  |  | pexiganan | 0.6/0.7<sup>§</sup> |
| $MIC$ | Dose of antimicrobials for which the net growth rate is 0 | melittin | 0.48/1.83<sup>§</sup> |
|  |  | pexiganan | 0.54/2.14<sup>§</sup> |
| $c$ | Cost of resistance | | 0.5 |
| $B(t=0)$ | Population size at time t=0 | | $10^8$ |
| $t_{max}$ | Maximal run of simulation | | 7d |
| $s_N$ | Switching rate from N to P | melittin | 0.0012 |
|  |  | pexiganan | 0.00021 |
| $s_P$ | Switching rate from P to N | melittin | 2.26/0.28<sup>§</sup> |
|  |  | pexiganan | 1.26/0.40<sup>§</sup> |

<sup>§</sup> Naïve/Primed

**Table S 6.** Strains and plasmids used in this work and their relevant phenotypes

| Strain | Relevant phenotype | Reference |
| --- | --- | --- |
| *E. coli* MG1655 WT | F<sup>-</sup> λ<sup>-</sup> *ilvG*<sup>-</sup> *rfb-50 rph-1* | Lab stock |
| *E. coli* MG1655 *csgA*::Kn | *csgA*::Kn F<sup>-</sup> λ<sup>-</sup> *ilvG*<sup>-</sup> *rfb-50 rph-1* | this work |
| *E. coli* MG1655 *csgA*::scar | *csgA*::scar* F<sup>-</sup> λ<sup>-</sup> *ilvG*<sup>-</sup> *rfb-50 rph-1* | this work |
| *E. coli* MG1655 *wza*::Kn | *wza*::Kn F<sup>-</sup> λ<sup>-</sup> *ilvG*<sup>-</sup> *rfb-50 rph-1* | this work |
| *E. coli* MG1655 *wza*::scar* | *wza*::scar* F<sup>-</sup> λ<sup>-</sup> *ilvG*<sup>-</sup> *rfb-50 rph-1* | this work |
| *E. coli* BW25113 *csgA*::Kn | *csgA*::Kn *lacI*<sup>+</sup>*rrnB*<sub>T14</sub> Δ*lacZ*<sub>WJ16</sub> *hsdR514* Δ*araBAD*<sub>AH33</sub> Δ*rhaBAD*<sub>LD78</sub> *rph-1* Δ(*araB-D*)567 Δ(*rhaD-B*)568 Δ*lacZ*4787(::*rrnB-3*) *hsdR514 rph-1* | Baba et al., 2006 |
| *E. coli* BW25113 *wza*::Kn | *wza*::Kn *lacI*<sup>+</sup>*rrnB*<sub>T14</sub> Δ*lacZ*<sub>WJ16</sub> *hsdR514* Δ*araBAD*<sub>AH33</sub> Δ*rhaBAD*<sub>LD78</sub> *rph-1* Δ(*araB-D*)567 Δ(*rhaD-B*)568 Δ*lacZ*4787(::*rrnB-3*) *hsdR514 rph-1* | Baba et al., 2006 |
| Plasmids | Relevant phenotype | Reference |
| pKD46 | oriR101 w/repA101ts Amp<sup>R</sup> | Datsenko and Wanner, 2000 |
| pCP20 | oriR101 w/repA101ts Amp<sup>R</sup> | Datsenko and Wanner, 2000 |

\* asterisk indicates the removal of the antibiotic resistant cassette kanamycin (Kn) by the action of the flippase expressed by the plasmid pCP20 (Datsenko and Wanner, 2000).

## Supplementary Figures

**Figure S 1.**
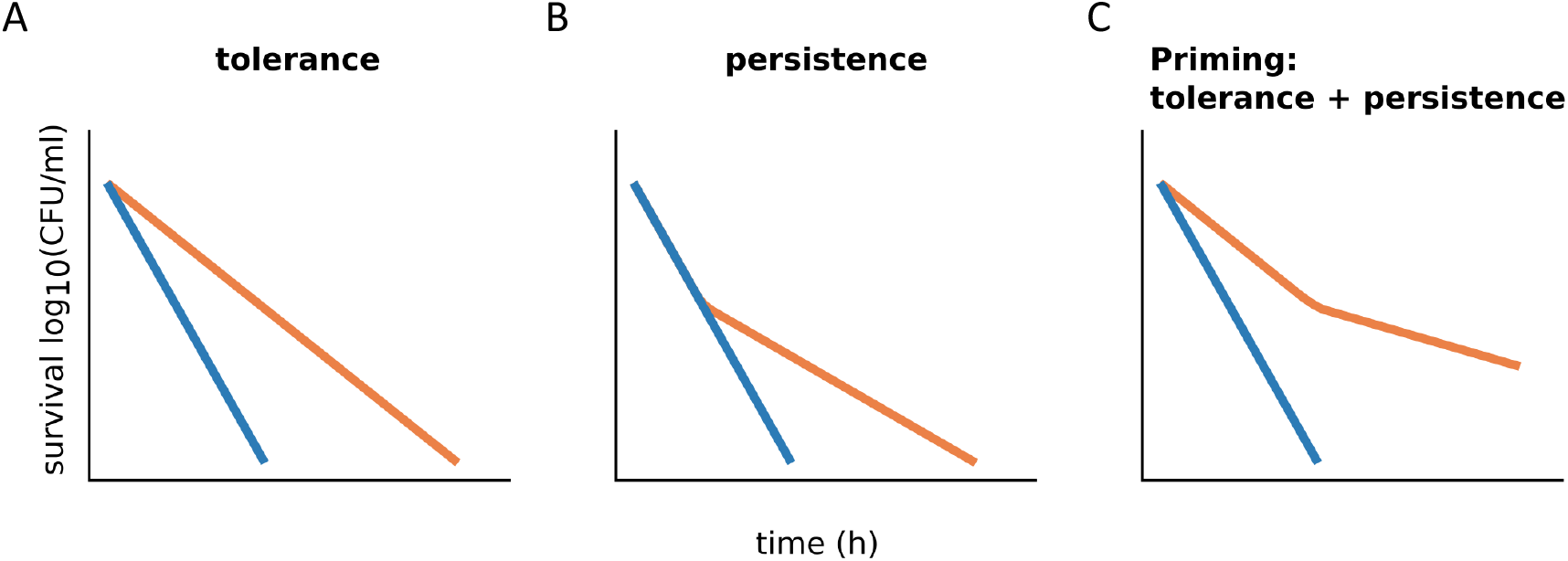
Bacterial tolerance and persistence change the shape of time-kill curves (figure adapted from Brauner et al. (*44*). (A) Tolerance is the ability of bacteria to longer survive exposure to antimicrobials due to decrease in susceptibility. Tolerance is quantified as increase in the slope of the time kill curve. (B) Persistence is the phenomenon of a subpopulation being less susceptible to the antimicrobial than the rest of the population. A persistent subpopulation manifests as a biphasic decline of bacterial population when exposed to lethal concentrations of antimicrobials. Here, the population consists predominantly of the less susceptible persistent subpopulation. (C) Together, tolerance and persistence result in biphasic time-kill curves with decreased susceptibility in the first phase.

**Figure S 2.**
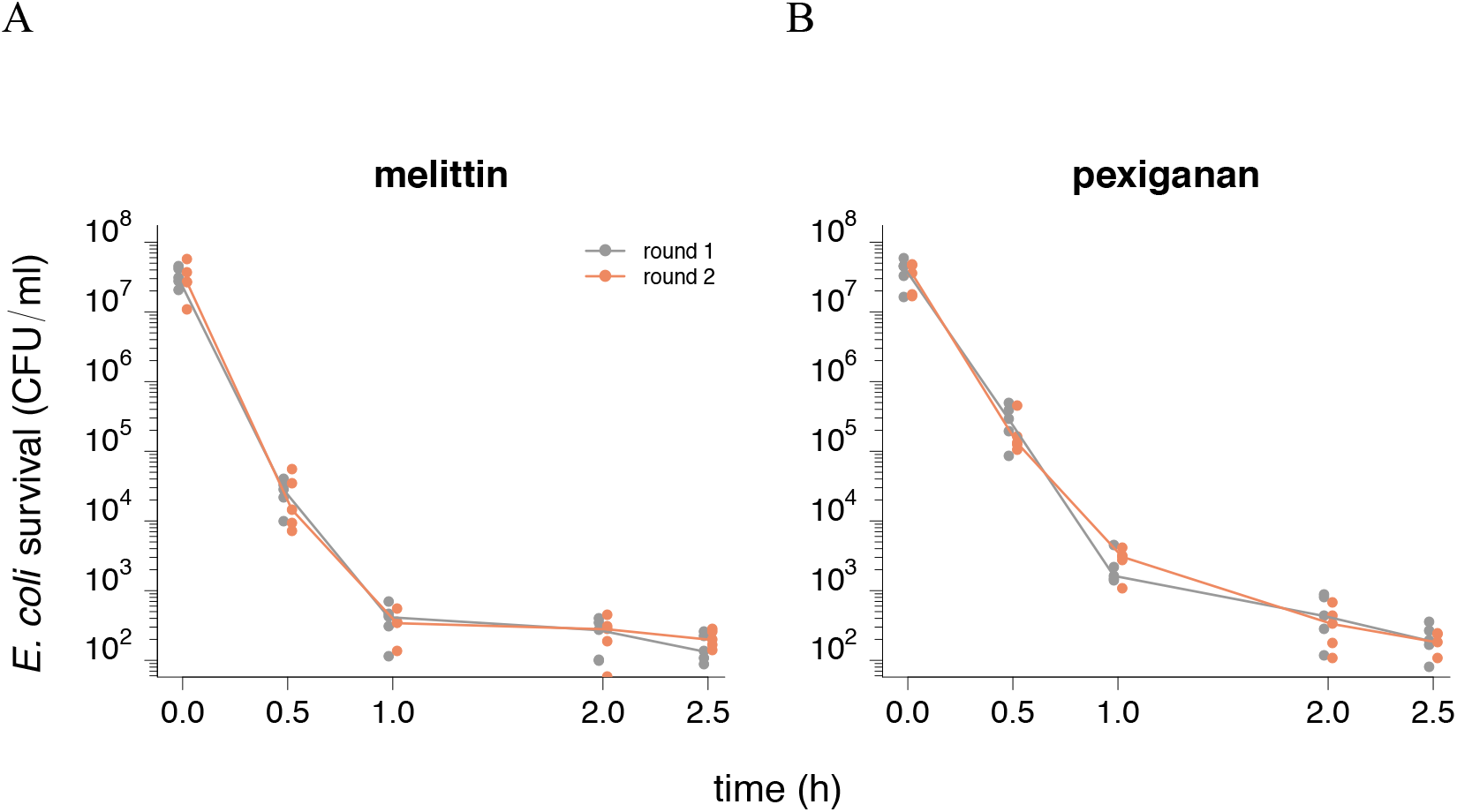
A possible explanation for a nonlinear decline in populations would be a decrease in active AMP concentration over time due to degradation (*45*, *46*), We tested this alternative explanation. As in Figure 1 (main text), we measured the population dynamics of bacteria exposed to 10 × MIC (round 1). At the end of the experiment, we sampled the supernatant. In round 2, a fresh bacterial population was exposed to the sampled supernatant. (A) and (B) show survival data of both rounds for melittin and pexiganan, respectively. The trend line depicts the median of the population size at each time-point. We tested differences in population size over time and differences between round one and two for each antimicrobial peptide with an ANOVA and adjusted with the Bonferroni method for multiple testing. For both, melittin and pexiganan, the population size changed significantly over time, while the differences in population size between round one and two and the interaction between time point and round were not significant (p <0.05).

**Figure S 3:**
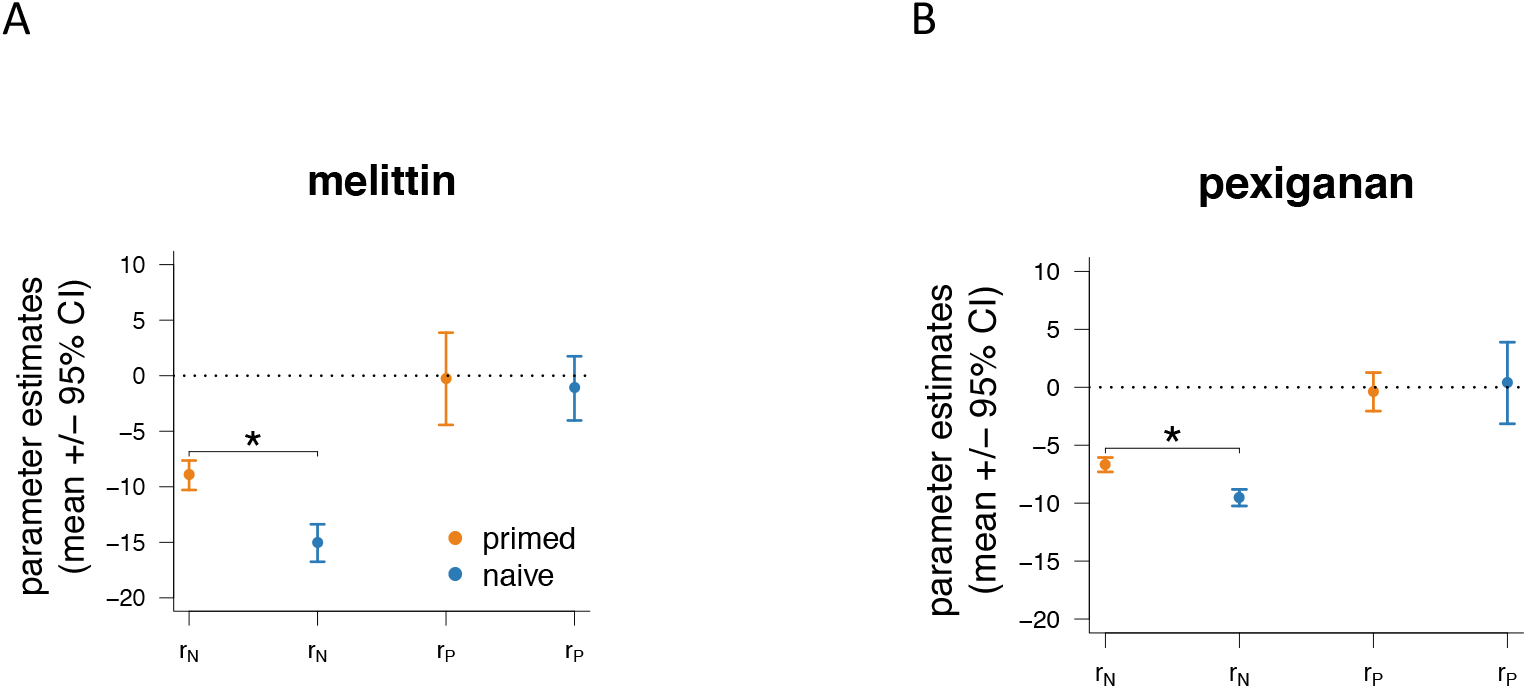
Net growth rates resulting from the fit of the two-state model with 4 free parameters. For both (A) melittin and (B) pexiganan, *r_P_* is not significantly different from 0. Significant differences between naive and primed treatment are indicated with asterisks.

**Figure S 4.**
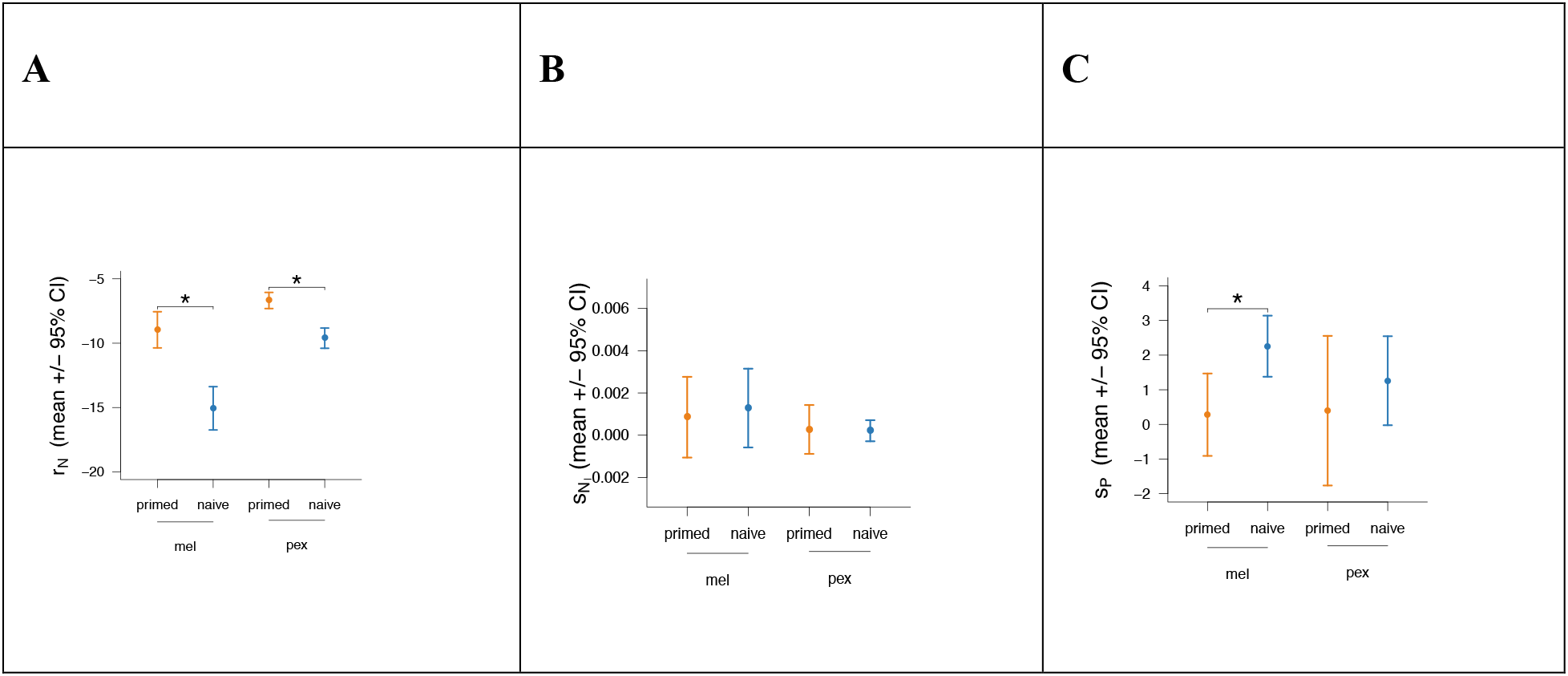
Parameter (A) r_N_, (B) s_N_, and (C) s_P_ of the two-state model fitted to the data of primed (orange) and naïve (blue) bacteria. The parameter *r_P_* was set to 0. Significant differences between naïve (blue) and primed (orange) parameter values are indicated with asterisks. For parameter values, see Table S 5.

**Figure S 5.**
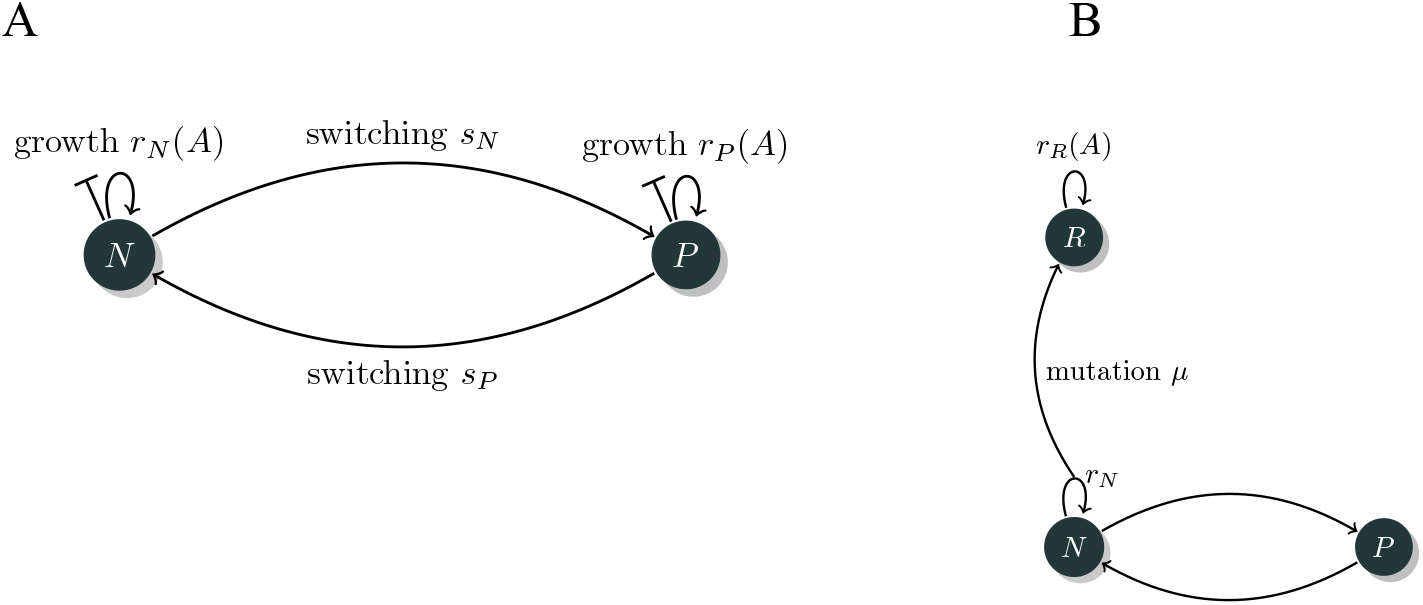
Diagrammatic representation of (A) the two-state model (*17*) (see also Fig. 5A) and (B) our previously developed framework (*38*), which we extended by a persistent class according to *(17)*. Here, bacteria that replicate mutate with a mutation rate **μ** to a more resistant subpopulation R.

**Figure S 6.**
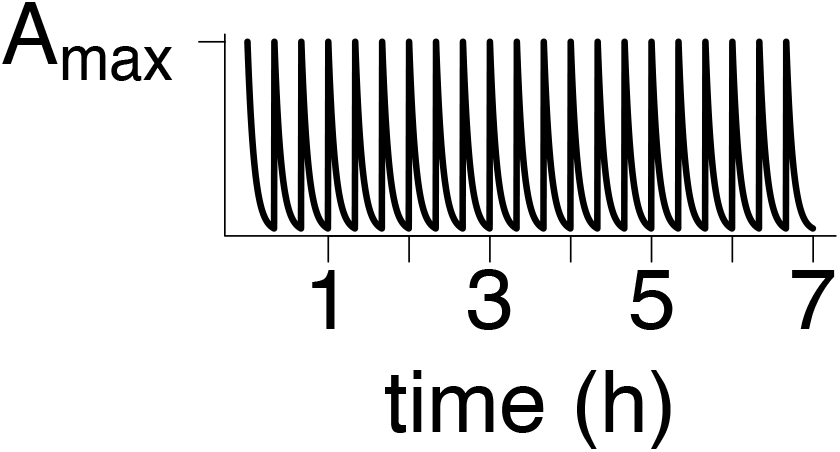
Pharmacokinetic profile used in our stochastic simulations.

**Figure S 7.**
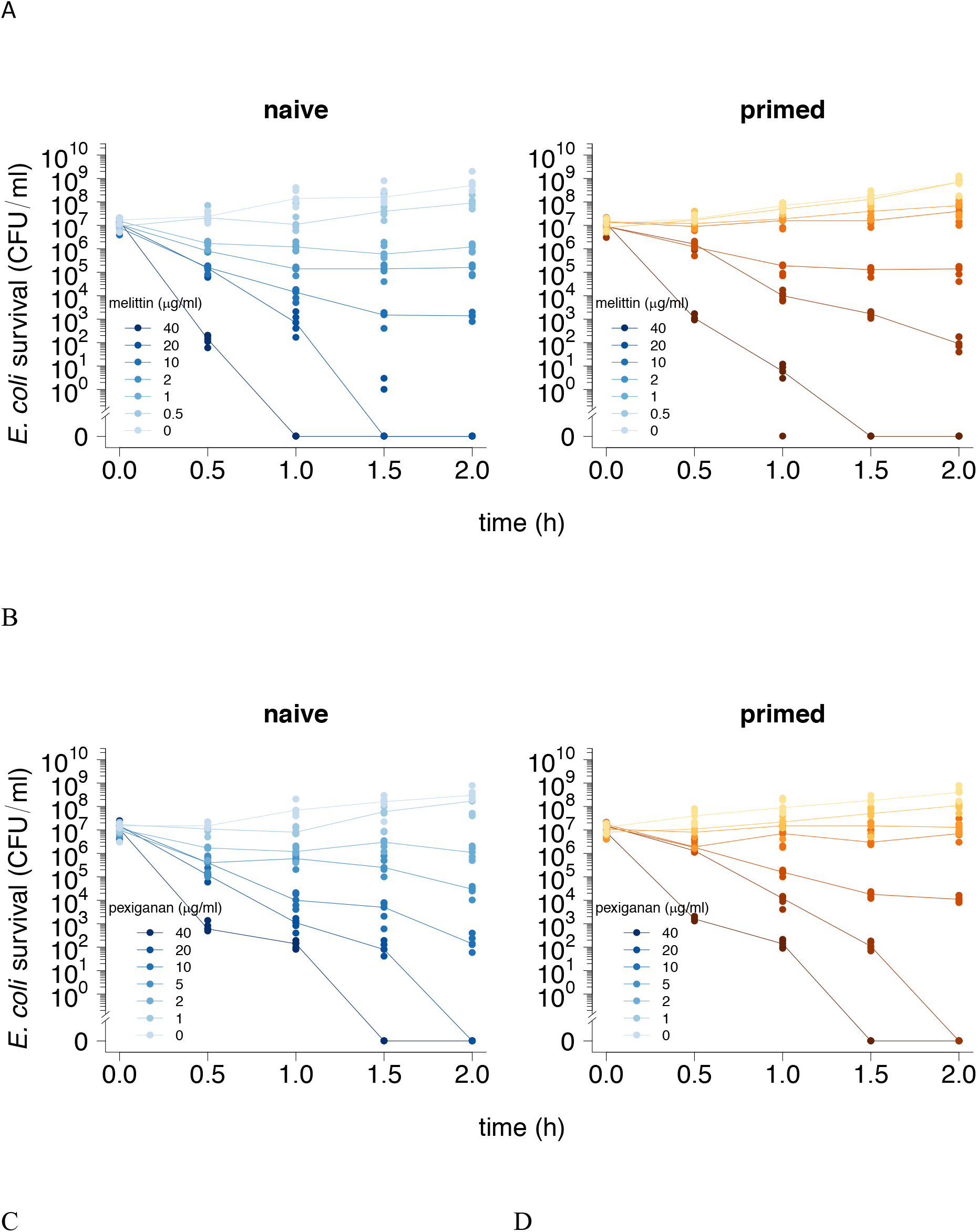
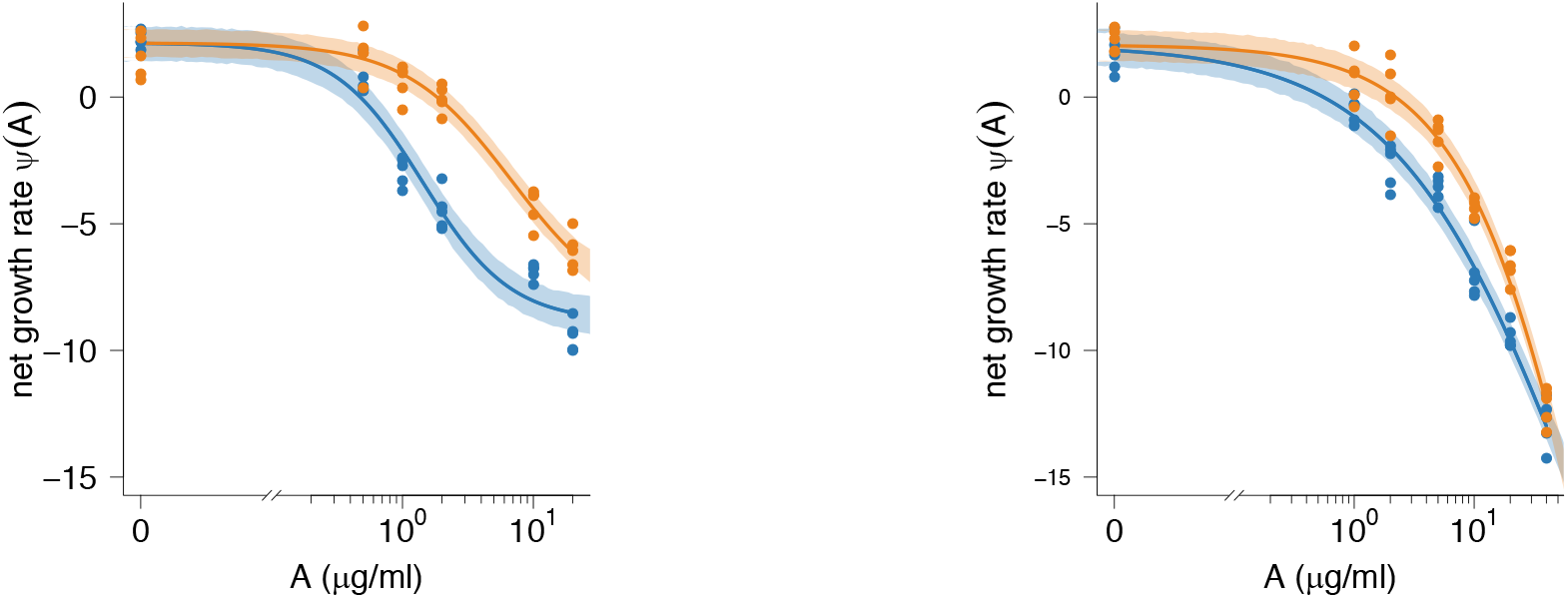
Time-kill curves and PD curves of bacterial population dynamics exposed to AMPs. Time-kill experiments in which naïve (blue) and primed (orange) bacteria were exposed to (A) melittin and (B) pexiganan. The AMP dose used in each time-kill experiment is listed in the legend in each plot. PD functions of melittin (C) and pexiganan (D) fitted to the data in (A) and (B), respectively. Note that we excluded data points of experiments in which naïve bacteria were exposed to melittin (40 μg/ml) from the analysis to ensure the best fit. Parameter values are listed in table S5.

**Figure S 8.**
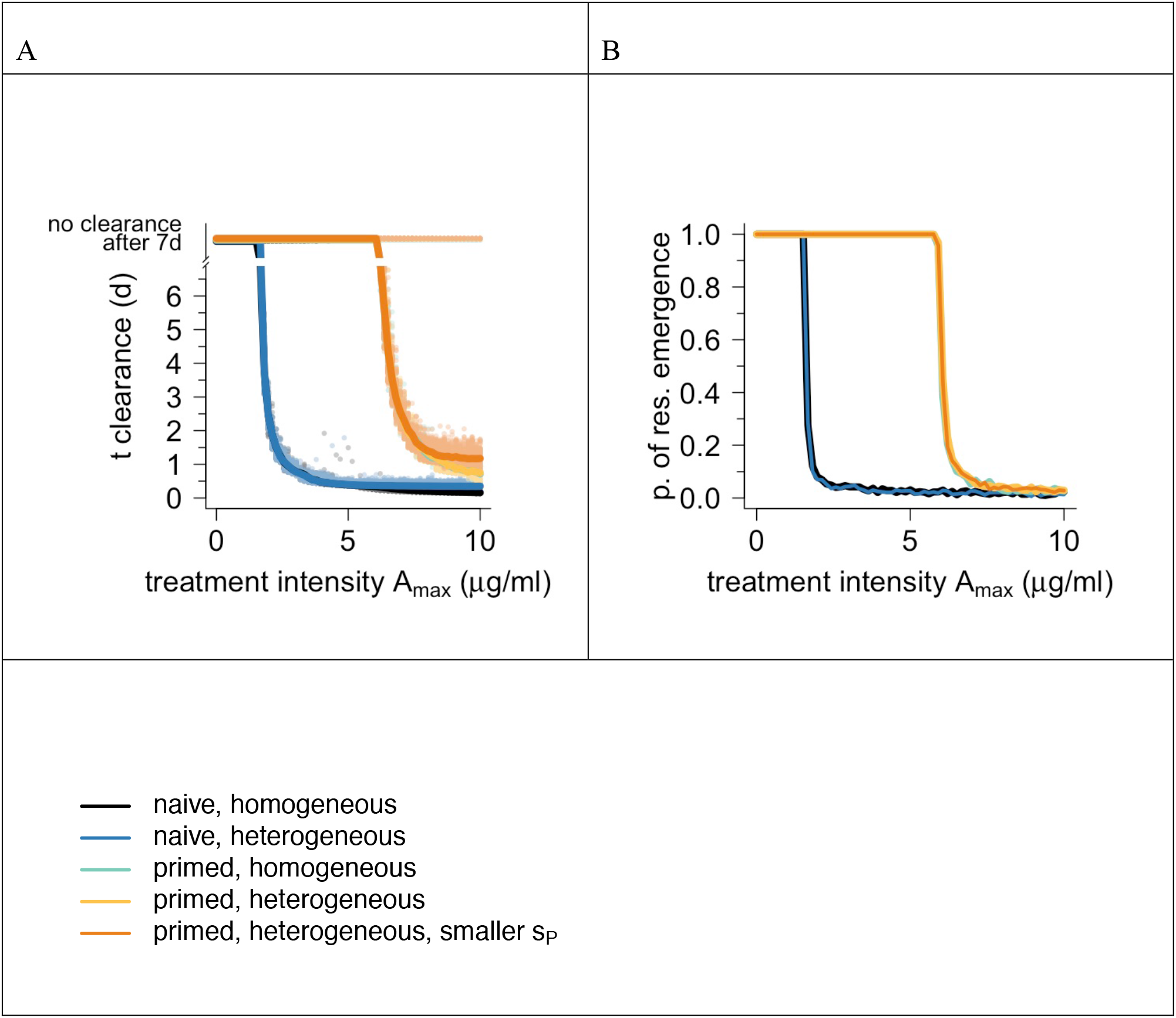
Time until clearance and probability of resistance evolution for bacteria exposed to pexiganan. In (A) each dot is an individual run.

**Figure S 9.**
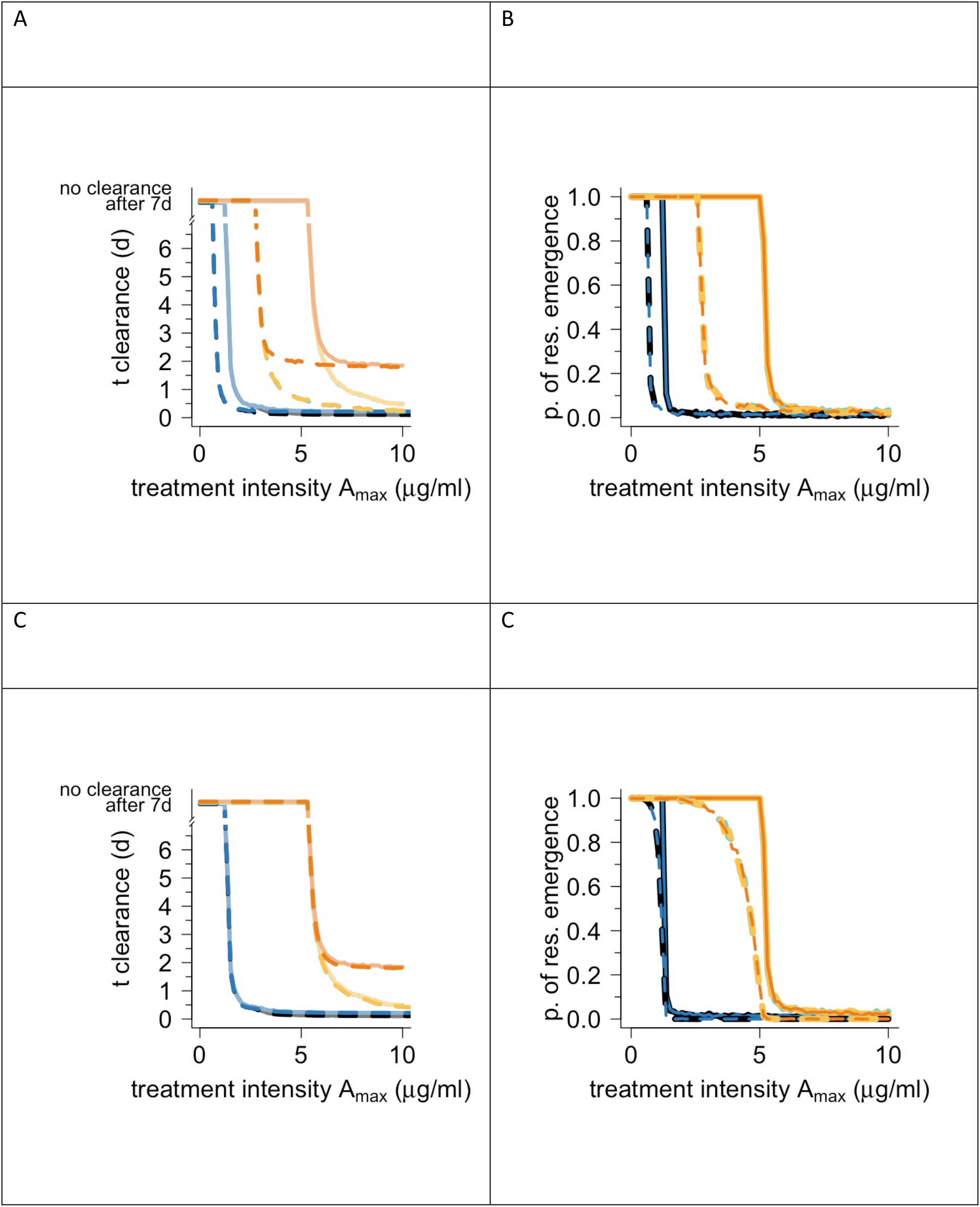
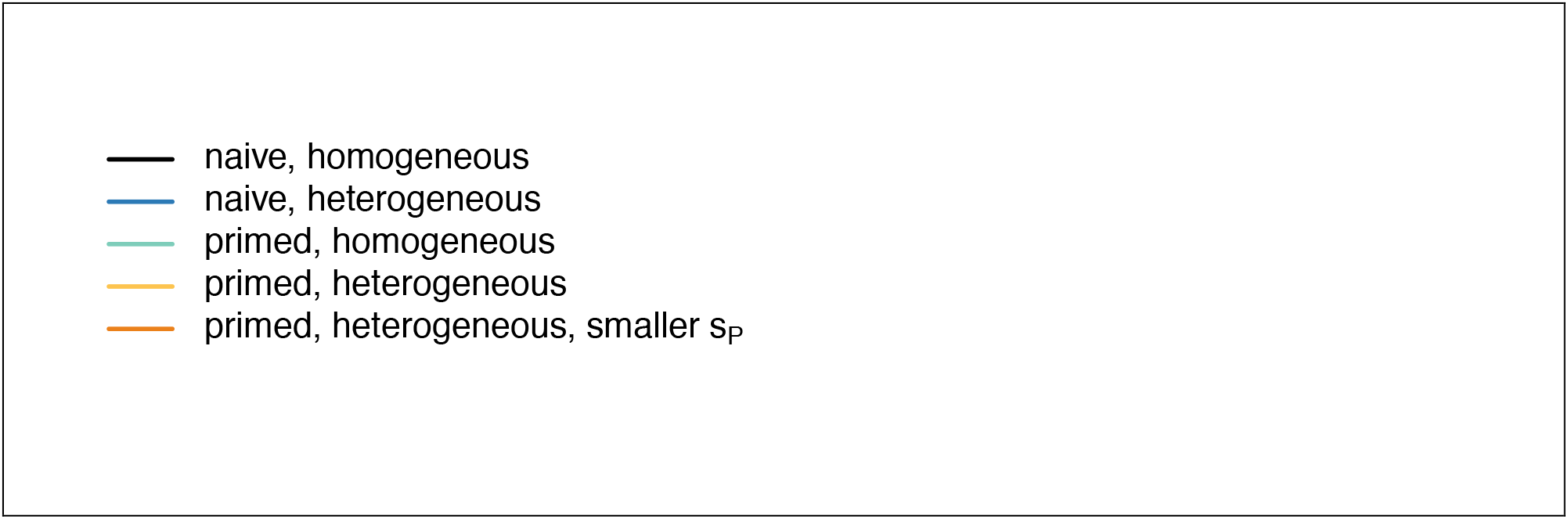
Predictions for different parameter values. (A) Time until clearance and (B) probability of resistance evolution is affected by the AMP decay rate k (dashed lines: k = 0.1, solid lines: k = 0.3). In (C) and (D) the mutation rate is varied (dashed lines: μ = 10−11, solid lines: μ = 10−9). All simulations are based on melittin data.

**Figure S 10.**
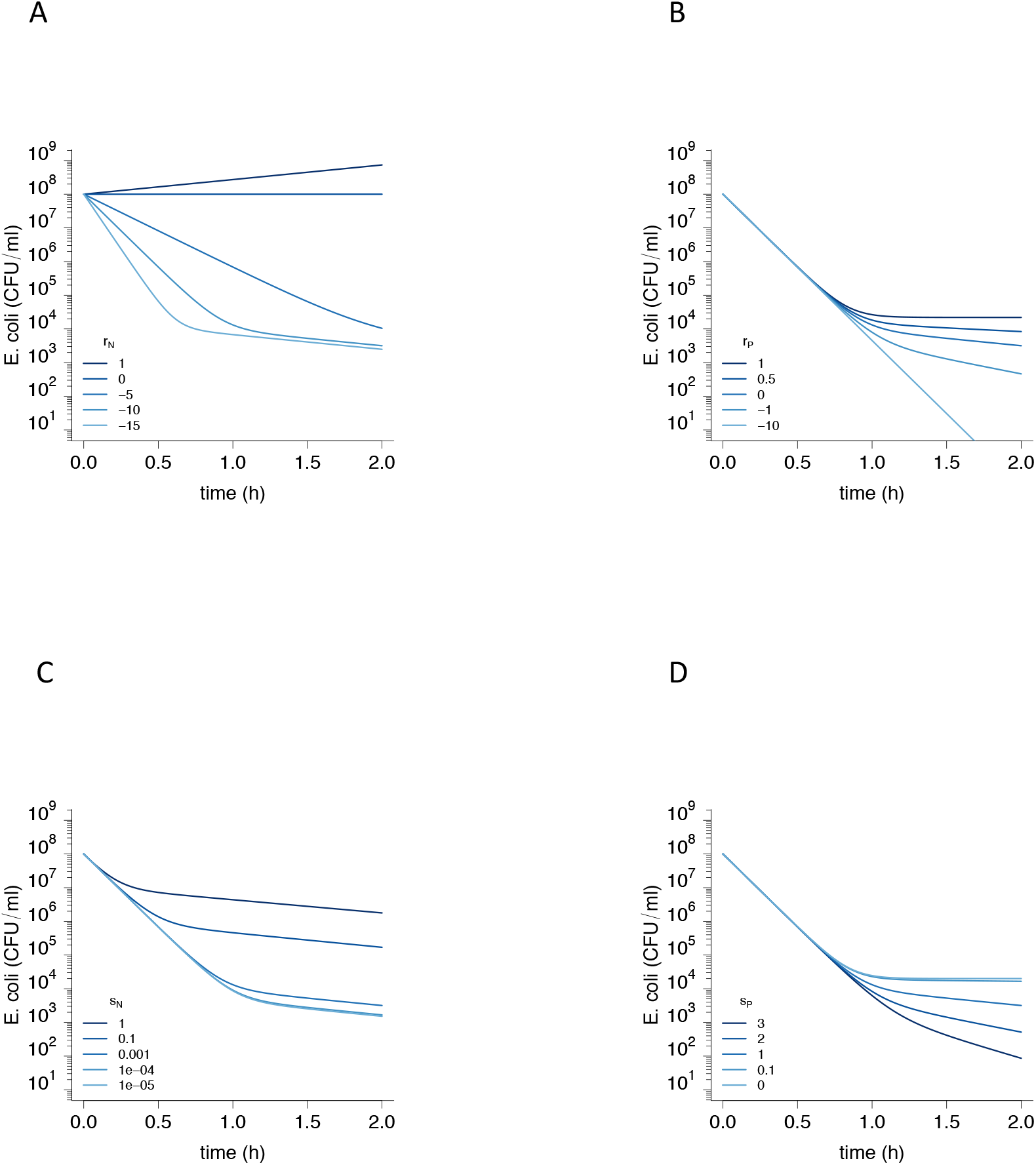
Two-state model predicts biphasic decline depending on the model parameter values. If not varied, *r_N_* = −10, *r_P_* = 0, *s_N_* = 0.001, and *s_P_* = 1. Tolerance, i.e. the slope *m_1_* is mainly influenced by *r_N_*, while the levels of persistence is influenced by all four parameters.

**Figure S 11.**
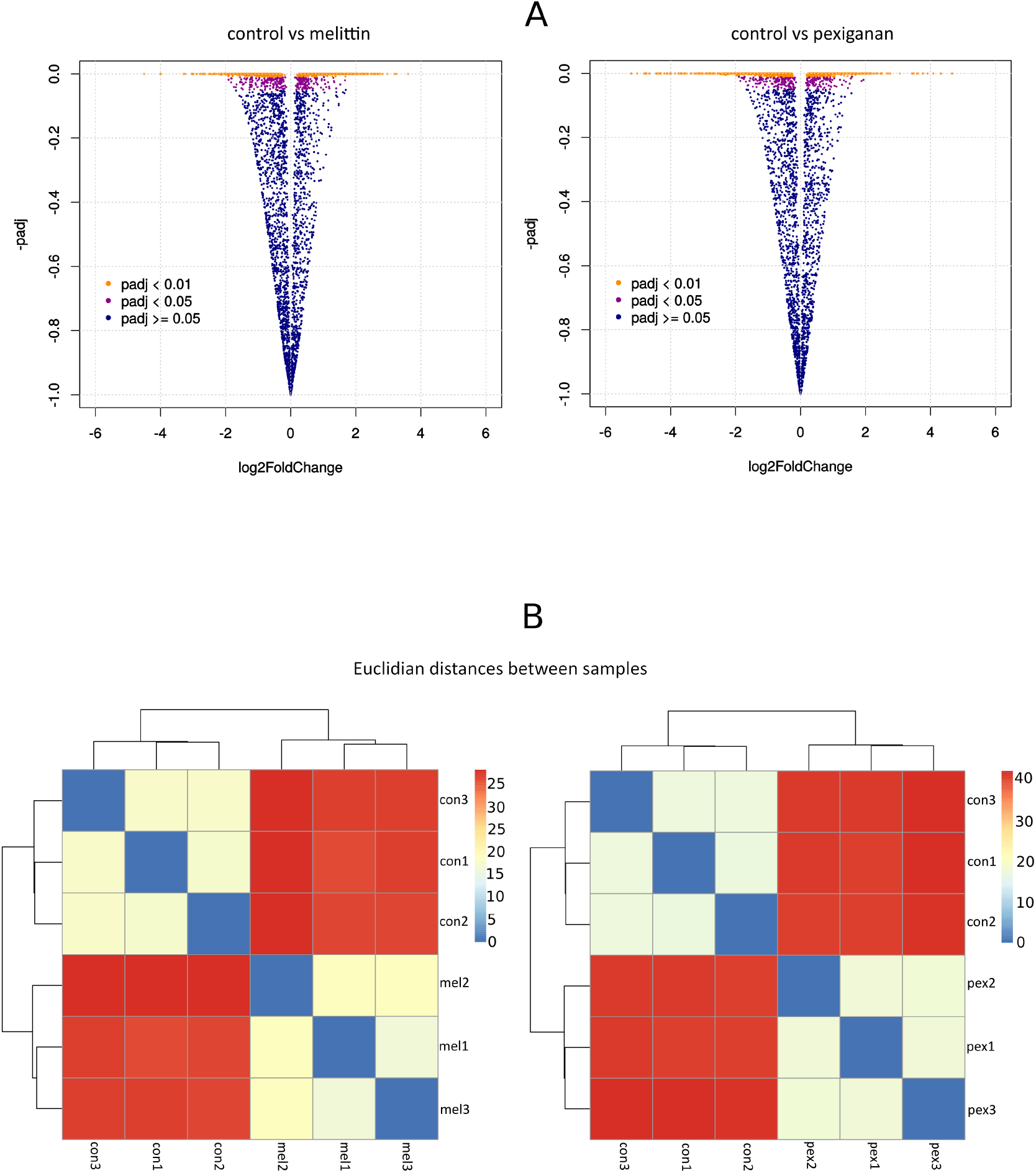
Quality control of RNA sequencing by evaluating symmetry and distribution of the transcriptome counts, volcano plots showing different degrees of significance (A) and assessing dissimilarities of sample-based Euclidian hierarchical clustering for cells treated with priming concentrations of melitting and pexiganan (B). Datasets are based on RNAseq of three independent biological replicas.

**Figure S 12.**
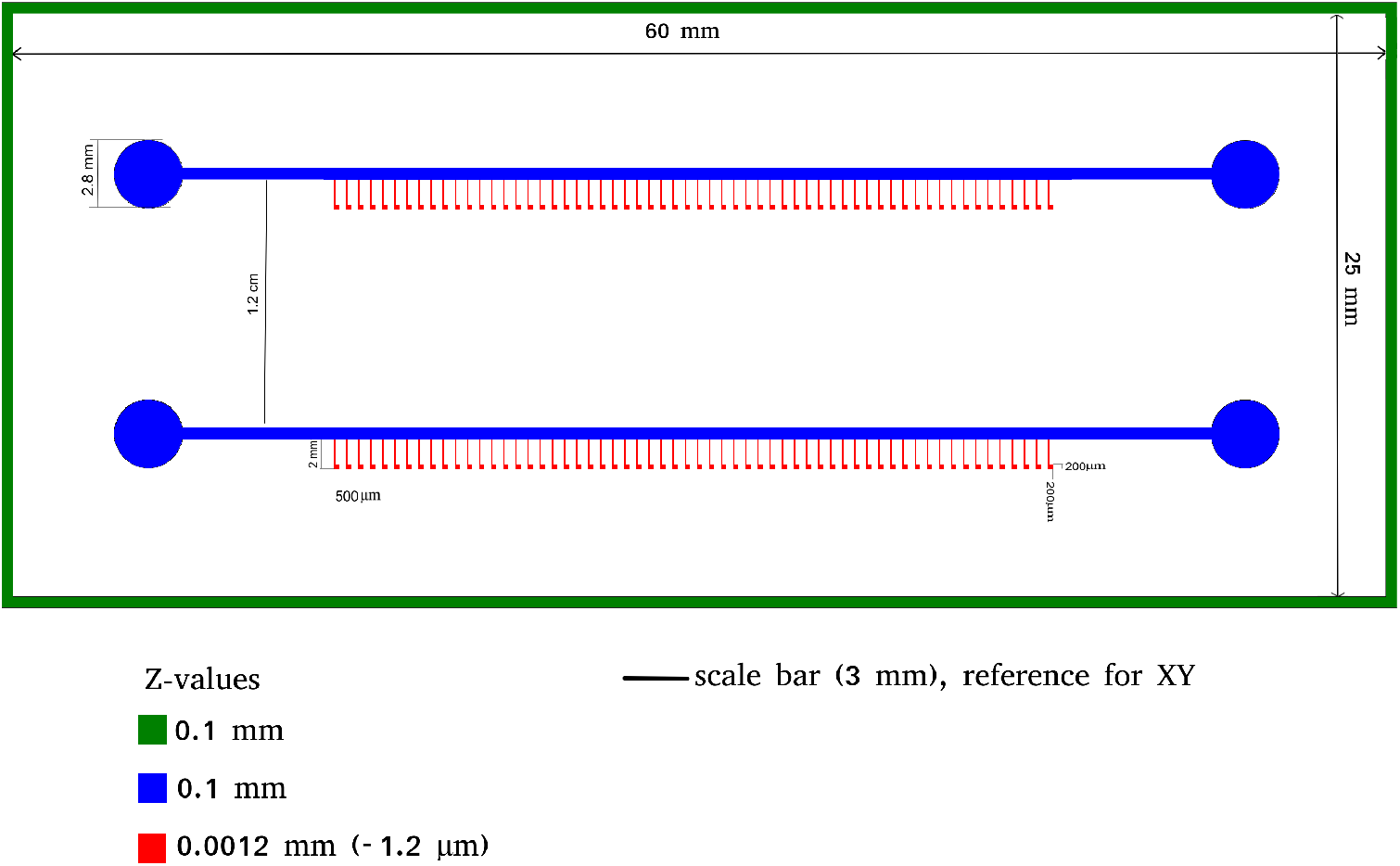
Microfluidic device design used for live imaging of priming and killing of *E. coli* MG1655 by AMPs. Z-values refers to the depth of chip features while X and Y represent the dimensions of 2D axes. Each chamber square compartment was designed to be similar in size (200 μm) to a microscope field with a magnification of 1000X.

